# Rewiring Mitochondrial Phosphatidylethanolamine Metabolism Identifies New and Unaccounted Trafficking Steps

**DOI:** 10.64898/2026.04.22.720193

**Authors:** Rashima Prem, Erica Avery, Juliana M. Marquez, Chi Xie, Steven M. Claypool

## Abstract

The distinct compositions of the two mitochondrial membranes are generated through a combination of phospholipids that mitochondria can make and those they take; both processes depend on a series of distinct lipid trafficking steps. Mitochondria make phosphatidylethanolamine (PE) through the action of the phosphatidylserine decarboxylase Psd1, an intermembrane space (IMS)-facing integral inner membrane (IM) protein. Psd1 has been proposed to act on its endoplasmic reticulum-derived substrate, phosphatidylserine (PS), after its transport to the mitochondrial outer membrane (OM) and either following its Ups2/Mdm35-mediated transport across the IMS to the IM or instead, on the IMS-side of the OM in a process enabled by the mitochondrial contact site and cristae organizing system (MICOS). Here, we implement a two-pronged Psd1 rewiring-based strategy predicted to either 1) circumvent the need for Ups2/Mdm35 and/or MICOS; or 2) selectively ablate the ability of Psd1 to work *in trans*. Our results with yeast harboring Psd1 targeted to the OM demonstrate that, with respect to mitochondrial PE production, Ups2/Mdm35 and MICOS indeed function within the IMS. Using yeast expressing a topologically inverted Psd1 chimera that faces the matrix, we identify previously unappreciated transbilayer lipid trafficking steps within the IM and show that Psd1 does not operate via a MICOS-organized *in trans* mechanism. Further, retained flux through inverted Psd1 when both Ups2/Mdm35 and MICOS are absent strongly implicates the existence of a major, yet presently unknown, mediator(s) of lipid movement across the IMS. Collectively, these data suggest a new model of how mitochondrial membrane diversity is established and maintained.

## Introduction

Biological membranes are chemically and compositionally diverse structures of foundational importance to life. Chemical diversity in lipids derives from their myriad chemical structures that are built from the combination of structurally and chemically distinct hydrophilic headgroups and acyl chains that differ in their length, saturation, hydroxylation and linkage [1, 2]. Membrane compositional diversity arises from the unique combination of different lipid components, and within cells, is found between the plasma membrane and organelles, and even between many membrane leaflets [3–5]. Thus, each cellular membrane contains lipids with unique individual properties and capacities surrounded by other lipids whose relative abundance is itself membrane-specific. How this combined diversity, which is critical for the assorted functions that occur in, on, and across a given biological membrane, is established and maintained is a fundamental question in cell biology presently riddled with knowledge gaps.

We approach the question of organellar lipid diversity through the lens of mitochondria, which have two chemically and compositionally diverse membranes [3, 5–8], the outer membrane (OM) and inner membrane (IM), contain biosynthetic pathways for two distinct glycerophospholipids, cardiolipin (CL) and phosphatidylethanolamine (PE), and depend on the acquisition of phospholipid products, and precursors needed for CL and PE synthesis, that they cannot themselves make [9, 10]. Unlike CL, which is only made in the mitochondrial IM where it largely remains, PE is formed via four distinct processes, three of which occur in the endoplasmic reticulum (ER) and two of which are considered major: the ER-resident CDP-ethanolamine and mitochondrial phosphatidylserine decarboxylase pathways [11]. The central figure in the mitochondrial PE pathway is phosphatidylserine decarboxylase 1 (Psd1; PISD in mammals), a self-cleaving heterodimer consisting of an IM-embedded, membrane-spanning β subunit noncovalently associated with a catalytically essential α subunit exposed to the intermembrane space (IMS) [12–14]. Psd1 decarboxylates phosphatidylserine (PS) to form PE. In the yeast *Saccharomyces cerevisiae* and mammals, PS is made in the ER [15, 16]; therefore, Psd1 must gain access to its substrate for mitochondrial PE synthesis to occur. Due to the intrinsic amphipathic character of phospholipids, this seemingly simple fact is achieved through a series of distinct steps. First, PS must be displaced from the ER membrane and move across the aqueous cytosol into the outer leaflet of the OM. In yeast, this transfer step is mediated at least in part by ERMES [17–19], a protein complex that physically bridges ER and mitochondrial membranes [20], whereas in mammals, the lipid transport proteins, ORP5 and ORP8, mediate the non-vesicular transport of PS from the ER to mitochondria [21]. Once in the outer leaflet of the OM, PS must be moved to the IMS-facing OM leaflet, a process partially mediated through the lipid scrambling activity of the abundant β-barrel OM protein, Por1 (VDAC1/2 in mammals) [22]. From here, two distinct mechanisms have been proposed to grant Psd1 access to PS: 1) transfer of PS from the OM to the IM by the Ups2/Mdm35 (PRELID3b/TRIAP1 in mammals) heterodimeric lipid transport proteins [23, 24]; and 2) decarboxylation of PS on the IMS-facing leaflet of the OM *in trans*, in a process that is enabled by the Mitochondrial Contact Site and Cristae Organizing System (MICOS complex) and does not require PS to cross the IMS [23]. While the contribution of Ups2/Mdm35 to Psd1-mediated PE production is strongly supported by a bevy of *in vitro* and *in vivo* studies across model systems [23–30], the putative MICOS-orchestrated *in trans* mode has not been rigorously tested in the decade since it was first described [23]. The lack of such vetting is an important oversight since the *in trans* model hinges on the assumption that the kinetic delay in synthesis of PE and phosphatidylcholine (PC; made in the ER by PE trimethylation) in MICOS-null yeast (lacking 6 MICOS subunits) in an *in vivo* radiolabeled-serine pulse chase paradigm [16, 19, 29, 31–35] reflects the inability of Psd1 to function *in trans*, a capability only demonstrated in an elegant, reductionist liposome reconstitution system [23]. Another consideration is that it is presently unclear whether the IM-anchored Psd1 even has the reach to span the distance created by MICOS-mediated junctions between the IM and OM [36–39]. Therefore, it remains unresolved whether MICOS promotes Psd1-based PE production via an *in trans*, or some alternate, mechanism.

The untested status of the *in trans* model of Psd1 function is also notable given the almost systemic redundancy found throughout lipid metabolism. As mentioned, Psd1 is one of four PE biosynthetic pathways in cells. In addition to ERMES, Vps13-Mcp1 complexes also participate in phospholipid flux between the ER and mitochondria [17]. Psd1 still converts PS to PE in mitochondria lacking Por1 [22], indicating that other unidentified OM proteins also have scrambling activity in this membrane. Redundancy is also reflected in the ability of cells to adapt to the absence of a known step in lipid metabolism. For example, even though MICOS-null yeast exhibit a significant decrease in the rate of PE and PC synthesis, their steady state mitochondrial lipid profile mirrors that of wild type yeast [23]. This may reflect the ability of MICOS-null cells to detect and homeostatically respond to membrane perturbations through an array of emerging mechanisms [40–46]. Thus, redundancy exists even when the responsible players and their mechanisms have yet to be identified. Such insight is critical because it underscores that there is more to learn about uncharacterized pathways coordinating mitochondrial phospholipid flux.

A powerful strategy recently developed to test for new or redundant lipid trafficking steps and identify possible mediators involves reducing lipid biosynthetic redundancy and then retargeting lipid synthetic activity(ies) to other membrane compartments using chimeric proteins and their subcellular targeting information. This rewiring approach has helped identify the existence of lipid trafficking between the ER and peroxisomes [47], the ability of PS made in peroxisomes, lipid droplets, or the matrix side of the IM to gain access to Psd1 [48], the ability of PE to move across the IMS in both directions which suggested a new model of how IM phospholipid diversity is established [49], and as combined with genetic screens, novel transcriptional regulators and mediators of lipid trafficking steps between organelles [17]. Here, we have implemented a rewiring-based strategy to specifically test the MICOS-based *in trans* mechanism of Psd1 function. Our results demonstrate that Psd1 does not operate via a MICOS-organized *in trans* mechanism. Further, they strongly implicate the existence of a major and presently unknown mediator(s) of lipid transport across the IMS, recognition of which allows us to posit a new model of how mitochondrial IM and OM diversity is established and maintained.

## Results

### Psd1 on the OM obviates the need for Ups2-mediated PS trafficking

Ups2/Mdm35 live and work in the IMS and in yeast, have a significant, albeit non-essential role, in mitochondrial PE biosynthesis, particularly when the demand for mitochondrial energy is high [24]. MICOS contains 6 core subunits, four of which are integral to the IM, and is formed from two modules, the MIC60 and MIC10 subcomplexes, both of which have the ability to bend membranes [36–39, 50, 51]. In yeast, the MIC60 module includes Mic60, the founding MICOS subunit [52, 53], and its peripheral binding partner, Mic19 [54]. MICOS serves as both a structural determinant of cristae junctions and, through interactions between Mic60/Mic19 and OM-resident components including the Translocase of the Outer Membrane (TOM) and Sorting and Assembly Machinery (SAM) complexes, as a scaffold between the IM and the OM [36, 37, 39, 55]. Therefore, while the main MICOS complex is intimately associated with the IM, its associations span the IMS and connects to the OM.

We previously generated a chimeric Psd1 construct targeted to the OM, termed OM-Psd1 [56], which subsequently allowed us to interrogate the ability of PE made outside the IM to gain access to this compartment [49]. We reasoned that OM-Psd1, by encountering PS on the cytosolic side of the OM, would circumvent the need for either Ups2-mediated PS flux from the OM to the IM or MICOS-organized Psd1 conversion of PS in the OM. To ultimately test this, and ensure that any noted phenotypes are representative, we initially compared the expression and growth of three independent clones of *psd2*Δ*psd1*Δ yeast expressing either WT IM-localized Psd1 (called IM-Psd1 for short) or OM-Psd1, each with a 3XFLAG tag appended to its C-terminus to track the severance of Psd1 into its β (detected with Psd1 antiserum) and α (FLAG) subunits (Figure 1A). The untransformed *psd2*Δ*psd1*Δ parental strain, which requires supplemental ethanolamine to make PE by the ER-resident CDP-ethanolamine pathway when grown on synthetic defined (SD) media, served as a control for both blotting and growth. While all three IM-Psd1 clones were expressed at similar levels, the amount of OM-Psd1 clone 2 was higher than the other two OM-Psd1 clones (Figure 1B). Interestingly, this increased expression correlated with reduced growth of OM-Psd1 clone 2 on synthetic defined agar plates in the absence or presence of supplemental ethanolamine and regardless of carbon source (Figure 1C, D; glycolysis-feeding dextrose for SD and oxidative phosphorylation-requiring lactate for SDLac). Given this atypical growth defect, OM-Psd1 clone 2 was omitted from relative growth analyses compared to IM-Psd1 clones (Figure 1D). The other two OM-Psd1 clones grew similar to each other, as did each tested IM-Psd1 clone. On three of the tested plates, the conglomerate growth of OM-Psd1 yeast was slightly but significantly less than IM-Psd1.

**Figure 1.**
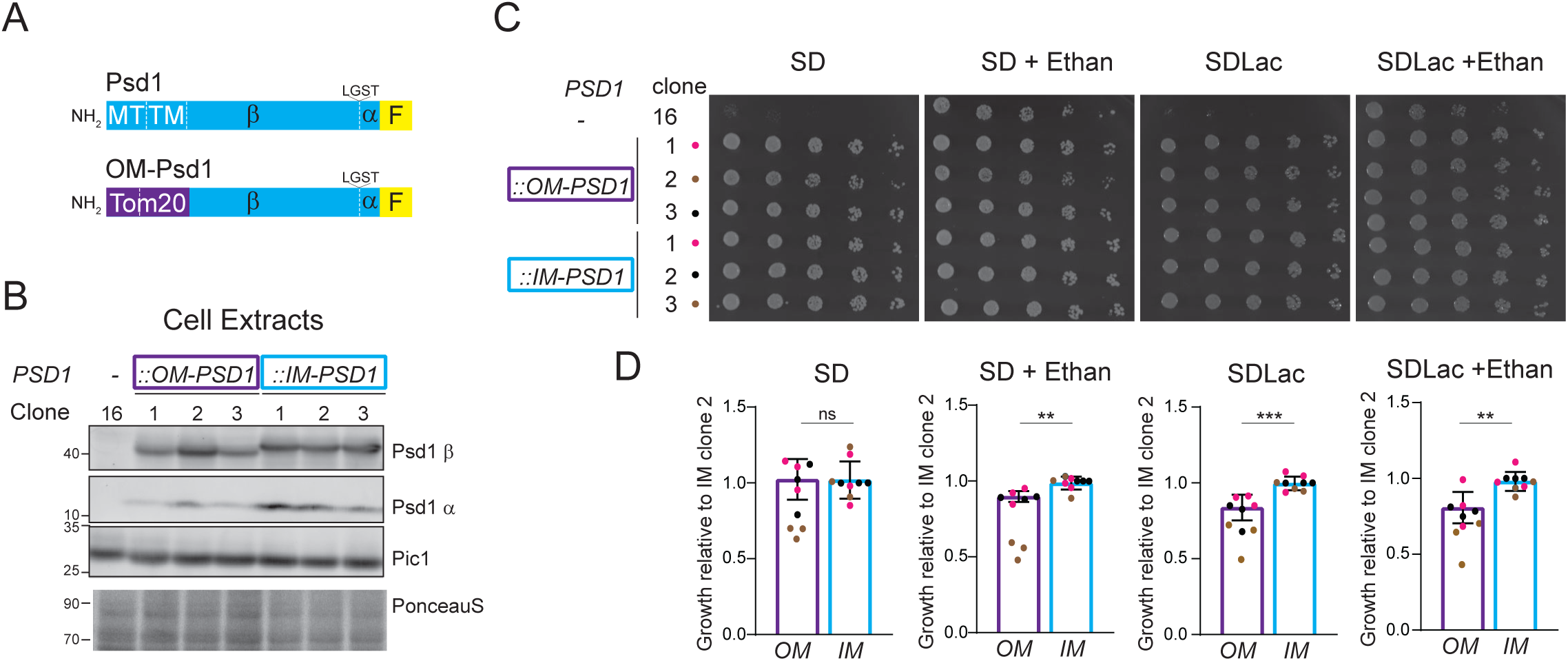
Identifying IM-Psd1 and OM-Psd1 clones with representative growth properties. (A) Cartoons of IM-Psd1 and OM-Psd1. Each construct contains a C-terminal 3XFLAG tag (F, yellow). The mitochondrial targeting (MT) and transmembrane (TM) domains of Psd1 were replaced by the equivalents from Tom20 (residues 1-100; purple). (B) Psd1 β and α subunits were detected by immunoblot in yeast cell extracts from 3 independent clones each for IM-Psd1 and OM-Psd1. Pic1 and total protein stain acted as loading controls. (C) The same strains were also spotted onto synthetic defined dextrose (SD) or lactate (SDLac) plates with or without 2mM ethanolamine and incubated at 30°C for 2 (SD) or 3 (SDLac) days. (D) Growth of each IM-Psd1 and OM-Psd1 clone relative to IM-Psd1 clone 2 was determined (mean ± SD for *n* = 9 biological replicates; individual clone-specific points are color-coded). Based on its anomalous growth behavior, relative growth data associated with OM-Psd1 clone 2 data was omitted for the growth comparisons between OM-Psd1 and IM-Psd1 yeast. Statistical differences (ns, *P* ≤ 0.05; 2 symbols, *P* ≤ 0.01; 3 symbols, *P* ≤ 0.001) between IM-Psd1 and OM-Psd1 were calculated by unpaired student t tests.

Using representative IM-Psd1 (clone 3) and OM-Psd1 (clone 1) parents, we generated a series of daughter strains that lacked Mic60 or Ups2 individually or in combination (Figure 2A,B). As expected, the steady state abundance of IM-Psd1 and OM-Psd1 was unchanged in the absence of Ups2, Mic60, or both (Figure 2B). To test potential phenotypic consequences stemming from the absence of Ups2, Mic60, or both in IM-Psd1 and OM-Psd1 yeast, growth was measured on synthetic media containing dextrose (SD) or lactate (SDLac), each as a function of ethanolamine supplementation (Figure 2C). Growth of both IM-Psd1 and OM-Psd1 yeast was largely unaffected by the absence of Ups2 (Figure 2C, D). As previously reported [23], the loss of Mic60 in IM-Psd1 yeast resulted in a growth defect on SD and SDLac which was partially restored when Ups2 was also absent (Figure 2D). The equivalent OM-Psd1 genotypes displayed the same growth pattern as their IM-Psd1 counterpart with one subtle exception: the significant difference in relative growth between *mic60*Δ and *mic60*Δ*ups2*Δ OM-Psd1 yeast was bridged upon addition of ethanolamine to SD. However, ethanolamine supplementation did not improve the overall growth of any IM-Psd1 or OM-Psd1 genotype, in stark contrast to the *psd2*Δ*psd1*Δ control. The lack of effect of ethanolamine suggests that PE is not limiting with respect to the growth of any of the tested IM-Psd1 or OM-Psd1 genotypes.

**Figure 2.**
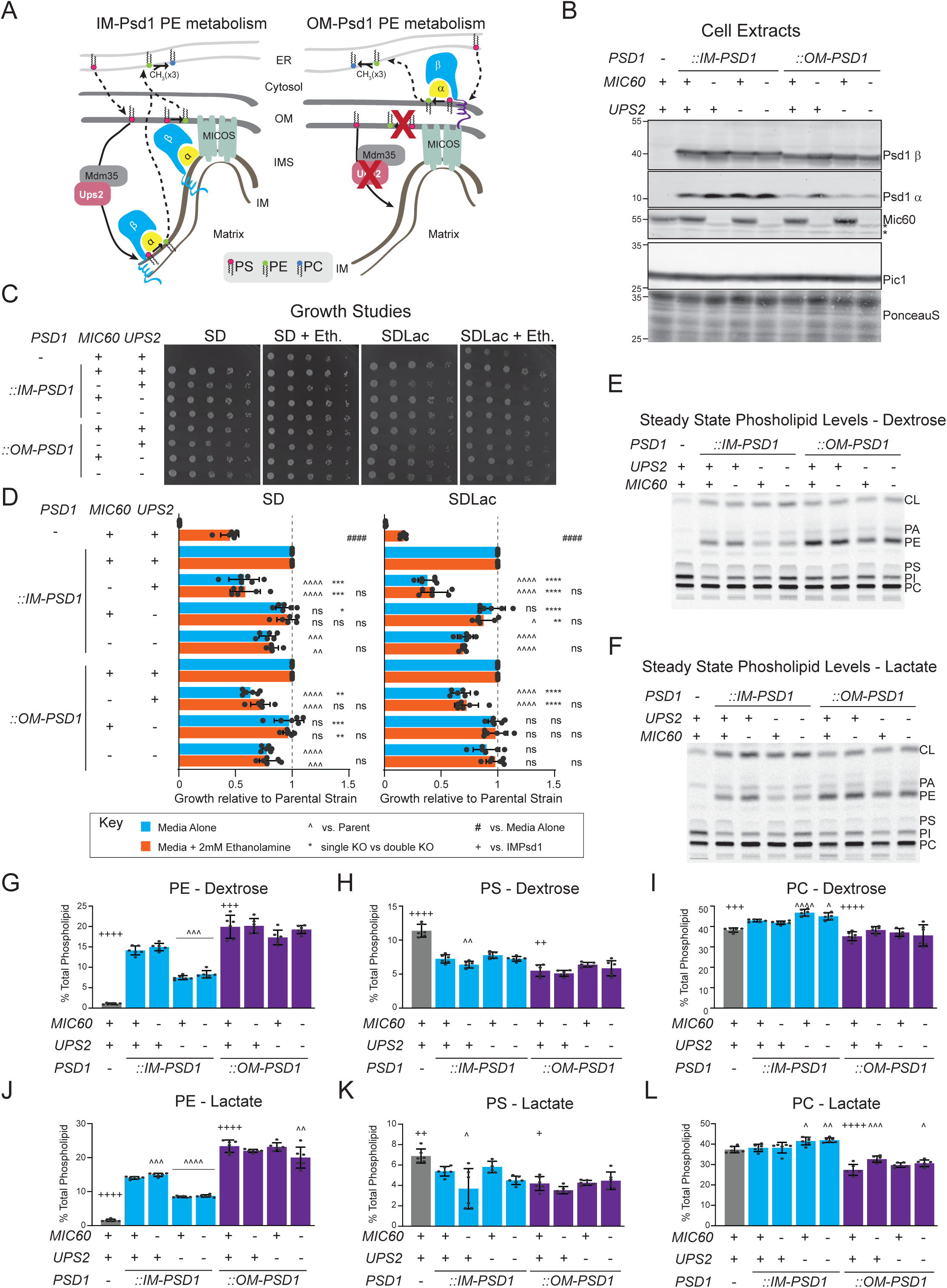
OM-Psd1 short-circuits the need for Ups2 or Mic60. (A) Cartoon outlining IM-Psd1 and OM-Psd1 PE metabolism highlighting the proposed roles for Ups2/Mdm35 and Mic60-containing MICOS. (B) Psd1 β and α subunits were detected by immunoblot in cell extracts from IM-Psd1 and OM-Psd1 yeast of the indicated genotypes. Pic1 and total protein stain served as loading controls. *, mark non-specific bands. (C) The listed strains were spotted onto SD and SDLac plates, each with or without 2mM ethanolamine and grown at 30°C for 2 (SD) or 3 (SDLac) days. PA, phosphatidic acid; PI, phosphatidylinositol. (D) Growth of each IM-Psd1 and OM-Psd1 daughter relative to their Ups2- and Mic60-proficient parent was determined; *psd2*Δ*psd1*Δ was compared to the IM-Psd1 parent (mean ± SD for *n* = 6 biological replicates). Statistical differences (ns, *P* > 0.05; 1 symbol, *P* ≤ 0.05; 2 symbols, *P* ≤ 0.01; 3 symbols, *P* ≤ 0.001; 4 symbols, *P* ≤ 0.0001) compared to the parent (^) or single versus double Ups2/Mic60 knockout (KO; *) were determined by one-way ANOVA with Sidak’s pairwise comparisons; differences as a function of ethanolamine were calculated by unpaired student t tests. Mitochondrial phospholipids were labeled overnight with ^14^C-acetate in indicated yeast grown in rich (E) dextrose or (F) lactate medium and resolved by TLC. Quantitation of mitochondrial PE (G, J), PS (H, K), and PC (I, L) amounts (mean ± SD for *n* = 6 biological replicates from 2 clones/genotype). Significant differences compared to the respective Ups2- and Mic60-proficient parent (^) or Ups2- and Mic60-proficient IM-Psd1 were determined by one-way ANOVA with Tukey’s multiple comparisons.

Next, we determined the steady state mitochondrial lipid profiles of the IM-Psd1 and OM-Psd1 panels following overnight growth in either rich dextrose (YPD) or lactate media, each spiked with ^14^C-acetate (Figure 2E, F and Figure S1). It was previously shown that respiratory growth of MICOS-deficient yeast is improved upon the additional loss of Ups2 or Psd1 [23]. This was taken as evidence that limiting PS flux into the IM and the resulting accumulation of PE could at least partially restore respiratory growth of MICOS-deficient yeast. Consistent with this possibility, the wildtype-like PE levels of IM-Psd1 yeast lacking Mic60 were reduced upon additional loss of Ups2 (Figure 2G, J); however, if anything, the slightly reduced PS amounts in the absence of Mic60 were restored when Ups2 was also gone (Figure 2H, K). Another carbon-source agnostic change noted for IM-Psd1 yeast lacking Ups2 alone or in combination with Mic60 was that PC levels were increased (Figure 2I, L). In OM-Psd1 yeast, PE levels were elevated as expected [49] and importantly, remained high in the absence of either Ups2 or Mic60, although PE levels were only slightly reduced upon their combined absence (Figure 2 G, J). These phospholipid results indicate that limiting the accumulation of PE is not responsible for the rescued growth of MICOS-deficient yeast upon additional loss of Ups2. Alternatively, the loss of Mic60 impacts growth of IM-Psd1 and OM-Psd1 yeast via different mechanisms, neither of which can be rescued with ethanolamine. Further, these findings are consistent with Ups2 and Mic60 functioning within the IMS and support the model that OM-Psd1 short-circuits the need for these factors with respect to the Psd1 pathway.

### Validating IM-anchored Psd1 with an inverted topology

To interrogate the MICOS-organized *in trans* model of Psd1 function, we adopted a strategy successfully used to direct PS synthase activity to the matrix side of the IM [48] to generate a Psd1 chimera embedded in the IM but with the opposite topology as IM-Psd1 (Figure. 3A). Like IM-Psd1 and OM-Psd1, inverted Psd1 (Inv-Psd1) was autocatalytically competent, producing both mature β and α subunits, and supported ethanolamine-free growth (Figure 3B, C). To test the IM topology of Inv-Psd1, we determined its protease sensitivity as a function of mitochondrial membrane integrity (Figure 3D). Both IM-Psd1 and Inv-Psd1 were protected from protease in intact mitochondria, unlike the OM-anchored Tom70. Upon formation of OM-ruptured mitoplasts (MP), both subunits of IM-Psd1 became sensitive to added protease. In contrast, the Inv-Psd1 α subunit remained protected from protease and its β subunit migrated as a smaller doublet, consistent with the degradation of its short IMS-exposed N-terminus.

**Figure 3.**
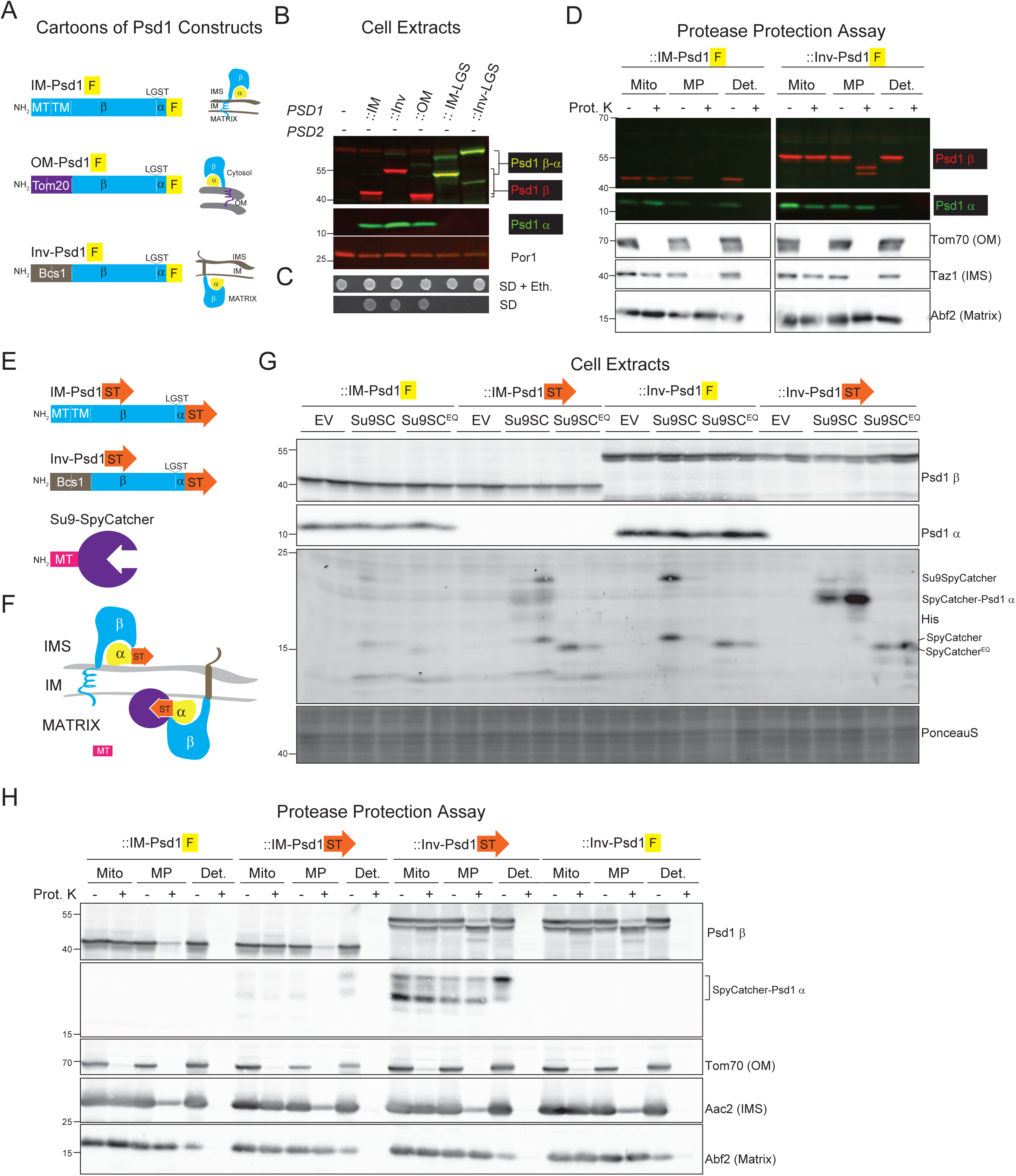
Inv-Psd1 is functional and topologically inverted in the IM. (A) Cartoons of IM-Psd1, OM-Psd1 and Inv-Psd1. Inv-Psd has a C-terminal 3XFLAG tag (F, yellow), like the other Psd1 chimeras, but the mitochondrial targeting (MT) and transmembrane (TM) domains of Psd1 were replaced by the Bcs1 equivalents (residues 1-128; brown). (B) Psd1 β and α subunits were detected by immunoblot in cell extracts from the designated yeast strains; IM-LGS and Inv-LGS are mutants of IM-Psd1 and Inv-Psd1 unable to execute autocatalysis. Por1 served as loading control. (C) The listed strains were spotted onto SD plates with or without 2mM ethanolamine and grown at 30°C for 3 days. (D) Protease accessibility assay in virgin mitochondria (Mito), OM-ruptured mitoplasts (MP), or detergent-extracted mitochondria (Det). After a 30min incubation in absence or presence of 0.1mg proteinase K (Prot. K), samples were harvested, separated by SDS-PAGE, and immunoblotted for the Psd1 β and α subunits. Tom70 (OM), Taz1 (IM facing IMS), and Abf2 served as compartment-specific controls. (E) Cartoons of IM-Psd1 and Inv-Psd1 with SpyTag2 added to their C-termini (ST, orange) and matrix-targeted Su9-Spycatcher002. (F) Illustration showing that an isopeptide bond between matrix-directed Spycatcher should form with Inv-Psd1ST, but not IM-Psd1ST. (G) Cell extracts from 2 clones per genotype were resolved by SDS-PAGE and immunoblotted for Psd1 (β subunit for all and α subunit for 3XFLAG-tagged specificity controls) and matrix-targeted SpyCatcher. PonceauS staining served as a loading control. The migration of different forms of SpyCatcher, including the adduct with the SpyTagged α subunit, are indicated on the right. (H) Submitochondrial localization of the Psd1 β subunit and SpyCatcher-Psd1 α subunit adduct determined by protease accessibility assay. Tom70 (OM), the N-terminus of Aac2 (IM facing IMS), and Abf2 (matrix) were markers of mitochondrial compartments.

Single-pass IM proteins with the N-terminus in the IMS and the C-terminus in the matrix are numerically few. As such, we sought to confirm the topology of Inv-Psd1 using a second, gain-of-signal strategy by exploiting the ability of the 14 amino acid SpyTag2 to form an isopeptide bond upon interacting with its complementary protein, SpyCatcher [57, 58] (Figure 3 E, F). Specifically, we replaced the 3XFLAG tag on IM-Psd1 and Inv-Psd1 with SpyTag2 and expressed these constructs in *psd2*Δ*psd1*Δ yeast together with either an empty vector (EV), matrix-targeted (Su9)-SpyCatcher, or as a control, Su9-SpyCatcherEQ (unable to form isopeptide bond with SpyTag2), with the latter two constructs also containing C-terminal His tags. Similarly transformed yeast expressing 3XFLAG-tagged IM-Psd1 or Inv-Psd1 served as an additional layer of control. Bands corresponding to unprocessed Su9-SpyCatcher and released SpyCatcher and SpyCatcherEQ were detected by immunoblot, with SpyCatcherEQ migrating slightly faster than its functional counterpart (Figure 3G). Importantly, a major His-reactive band that migrated at the expected size of a SpyCatcher-α subunit adduct (∼21kDa), was clearly detected in cell extracts from yeast expressing Inv-Psd1 harboring a SpyTag when a functional Su9-SpyCatcher was co-expressed. Although less abundant, a similar adduct was also detected for SpyTagged IM-Psd1. We postulated that the presence of such an adduct could reflect a chance encounter as Su9-SpyCatcher is passing through the IMS during its import into the matrix. Consistent with this notion, the SpyCatcher-α adduct detected for SpyTagged IM-Psd1 was sensitive to protease when the OM was ruptured, whereas the Inv-Psd1 equivalent remained protected and only became accessible to protease upon the inclusion of detergent to solubilize all membranes (Figure 3H). These results demonstrate that Inv-Psd1 is embedded in the inner mitochondrial membrane with its active site facing the matrix.

### Flux through Inv-Psd1 reveals new IM lipid trafficking steps

With the goal of developing a more physiologically relevant model, we implemented a knock-in (KI) strategy in which 3XFLAG-tagged IM-Psd1 or Inv-Psd1 were used to replace the functional *TRP1* allele occupying what had been the endogenous *PSD1* open reading frame in the *psd2*Δ*psd1*Δ strain (Figure 4A). Indeed, the steady state amounts of the β subunits from two independent clones of both IM-Psd1 and Inv-Psd1 were comparable to endogenous Psd1 detected in *psd2*Δ yeast (Figure 4B, D). As expected, the released α subunit was detected in each set of IM-Psd1 and Inv-Psd1 KI clones, although its steady state abundance was increased for Inv-Psd1 KI. While the basis for this increase is unclear, it could reflect a homeostatic response to making PE on the matrix side of the IM or simply reflect inherent variability in detecting small proteins by immunoblot (the 3XFLAG tagged α subunit is ∼12 kDa). Consistent with the separate detection of β and α subunits, knocked-in IM-Psd1 and Inv-Psd1 were functional based on their ability to grow without ethanolamine (Figure 4C). Like IM-Psd1 and the mitochondrial IM protein, Pic1, Inv-Psd1 co-sedimented with the mitochondria-enriched P13 fraction following differential centrifugation of yeast cell homogenates (Figure 4E). Finally, the differential protease sensitivity of the α and β subunits in OM-ruptured mitoplasts from Inv-Psd1 versus the IMS-facing IM-Psd1 confirmed that the business end of Inv-Psd1 resided in the matrix.

**Figure 4.**
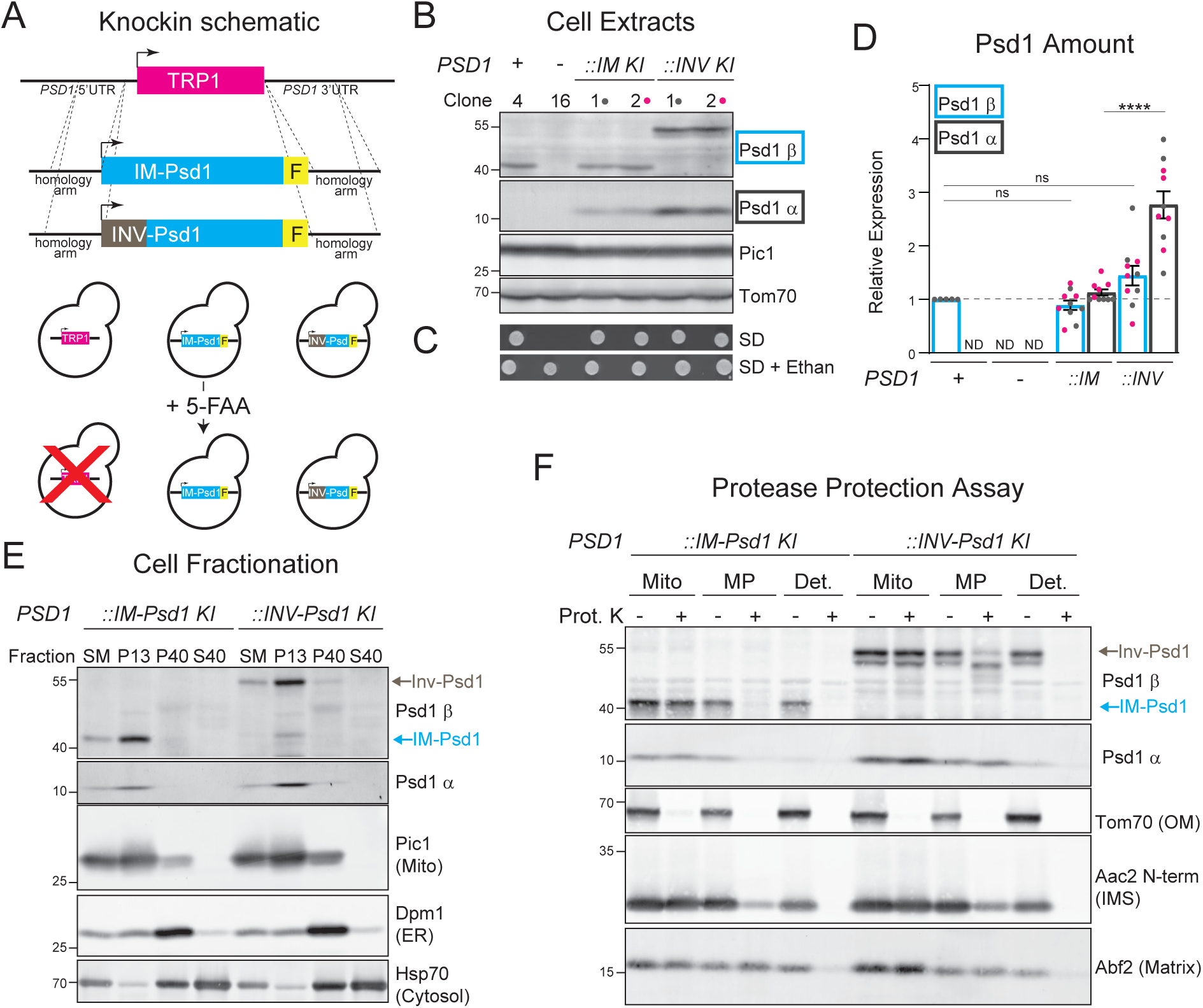
IM-Psd1 and Inv-Psd1 knock-in strain validation. (A) Illustration of IM-Psd1 and Inv-Psd1 knock-in and TRP1 counter-selection strategy. 5-FAA, 5-Fluoroanthranilic acid. (B) Psd1 β and α subunits were detected by immunoblot in yeast cell extracts from 2 independent knock-in clones each for IM-Psd1 and Inv-Psd1. Pic1 and Tom70 acted as loading controls. (C) The listed strains were spotted onto SD plates with or without 2mM ethanolamine and grown at 30°C for 2 days. (D) The amount of Psd1 β subunit for IM-Psd1 and Inv-Psd1 relative to endogenous Psd1 in *psd2*Δ yeast, and the amount of Psd1 α subunit in the knock-in clones relative to IM-Psd1 KI clone 1 were determined (mean ± SD for *n =* 10 biological replicates from 2 clones/Psd1 chimera; clone-specific data in pink or gray). (E) Fractions of lactate-grown IM-Psd1 and Inv-Psd1 knock-in yeast were harvested by differential centrifugation and equal protein amounts resolved by SDS-PAGE and immunoblotted for Psd1 β and α subunits and mitochondrial (Pic1), ER (Dpm1), and cytosolic (Hsp70) controls. SM, starting material; P13, pellet of 13,000 x *g*; P40, pellet of 40,000 x *g*; S40, supernatant of 40,000 x *g*. (F) Submitochondrial localization of Psd1 β and α subunits of IM-Psd1 and Inv-Psd1 determined by protease accessibility assay. Tom70 (OM), the N-terminus of Aac2 (IM facing IMS), and Abf2 (matrix) were mitochondrial compartment controls.

With the goal of comparing IM-Psd1 and Inv-Psd1 functionality *in vivo*, we initially ablated *DPL1* in both of the KI strains (2 clones each). Dpl1 is a minor source of ethanolamine phosphate derived from sphingolipid catabolism and can be used to produce PE by the CDP-ethanolamine pathway [59, 60]. Next, we compared IM-Psd1 and Inv-Psd1 activities in isolated mitochondria (Figure 5A, B). To do so, we adapted an established assay that measures the trafficking- and Psd1-dependent conversion of NBD-PS to NBD-PE [22, 24] by including TX-100 to grant equal access of substrate to both sides of the IM. By this approach, time-dependent accumulation of NBD-PE was indistinguishable between IM-Psd1 and Inv-Psd1 mitochondria. From this, we conclude that IM-Psd1 and Inv-Psd1 have the same intrinsic Psd activities. Finally, we compared flux through IM-Psd1 and Inv-Psd1 using an assay that tracks the sequential conversion of ^14^C-labeled PS to PE to PC (Figure 5C) [16, 19, 29, 31–35]. Upon addition of ^14^C-serine to cultures, it is rapidly incorporated into PS in the ER by the PS synthase, Cho1 [61]. These IM-Psd1 and Inv-Psd1 strains lack Psd2 and Dpl1 such that in the absence of these minor sources of ^14^C-serine derived PE, the subsequent accumulation of ^14^C-PE is solely Psd1-dependent and reflective of ^14^C-PS having navigated trafficking steps needed to become accessible to Psd1. Similarly, the appearance of ^14^C-PC indicates that PE has moved from its site of synthesis back to the ER, where the PE methyl transferases reside [62]. Notably, final access of PS to Inv-Psd1 requires transbilayer movement of PS to the matrix-facing IM leaflet; similarly, the first step after PE is made by Inv-Psd1 reports on the ability of PE to first flip to the IMS-side of the IM (Figure 5D). We therefore applied this ^14^C-serine pulse-chase paradigm and tracked the time-dependent transition of ^14^C-labeled PS to PE and finally to PC (Figure 5E, F). Compared to IM-Psd1, ^14^C-PS was depleted faster in Inv-Psd1 yeast and resulted in higher amounts of radiolabeled PE that persisted throughout the chase. Interestingly, the time-dependent increase in ^14^C-PC was the same for Inv-Psd1 as IM-Psd1. These results demonstrate that PS and PE can traverse IM leaflets in the manner needed for Inv-Psd1 to function in this circuit. Further, they suggest that the ability of PE to move from the matrix-side to the IMS-side of the IM is kinetically slower than PS movement in the opposite direction, resulting in higher PE amounts, but once exposed to the IMS, PE still traffics to the ER and is converted to PC just like for IM-Psd1.

**Figure 5.**
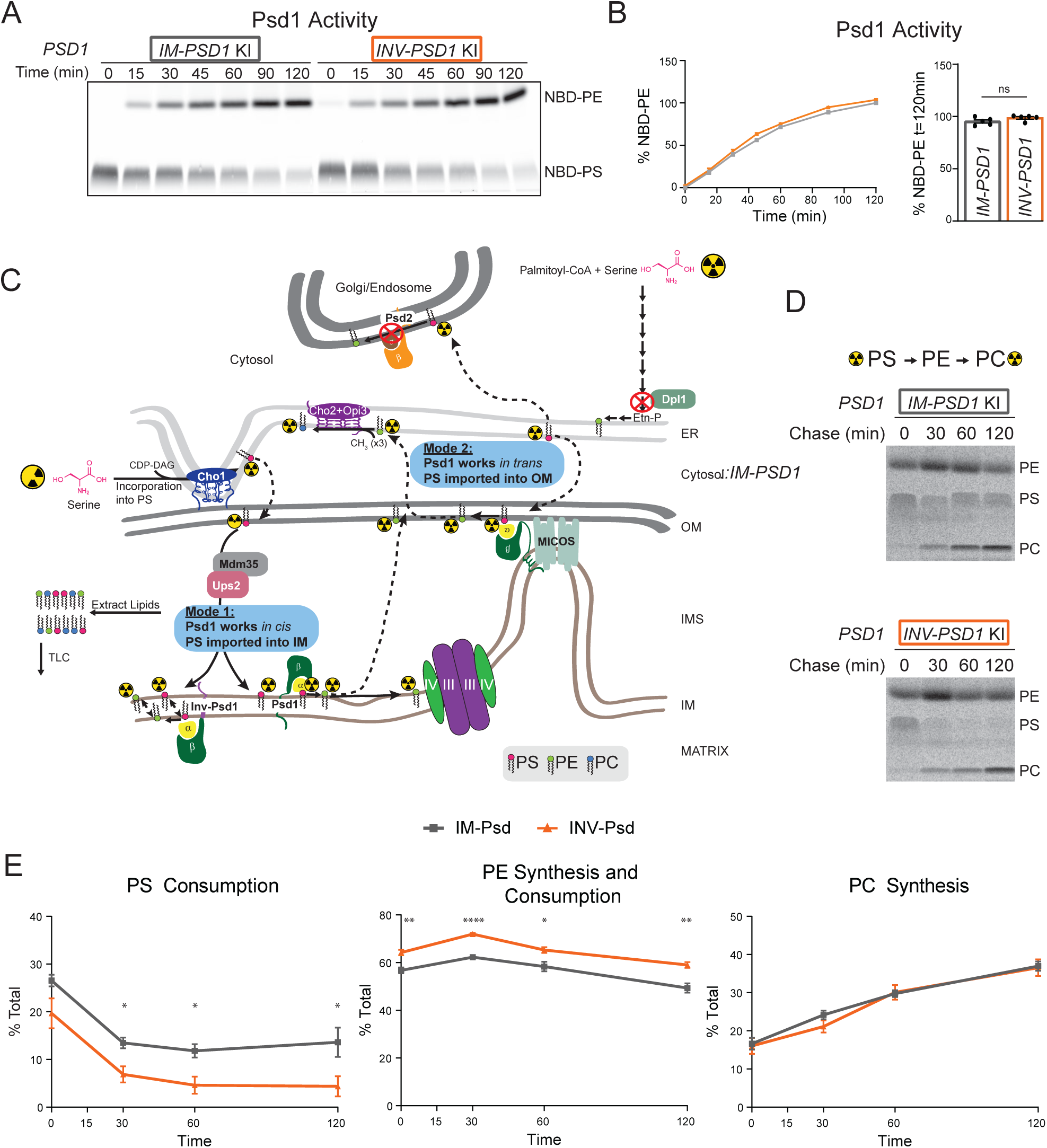
Flux through Inv-Psd1 is intact but slightly perturbed relative to IM-Psd1. (A) Time-dependent Psd1 activity in mitochondria from IM-Psd1 and Inv-Psd1 knock-in yeast was detected by decarboxylation of NBD-PS to NBD-PE following separation by TLC. (B) The time dependent accumulation of NBD-PE ((NBD-PE)/(NBD-PS + NBD-PE) x 100) and % NBD-PE (of total signal) at the final timepoint was calculated (mean ± SD for *n* = 5 biological replicates from 2 batches of mitochondria/genotype). (C) Illustration of Psd1-selective PE metabolism enforced by genetic disruption of minor pathways performed by Psd2 and Dpl1 using radioactive serine. (D) IM-Psd1 and Inv-Psd1 yeast were pulse-labeled with ^14^C(U)L-serine for 15min, washed, and then chased in SD with nonradioactive serine shaking at 30°C before removing aliquots at the indicated timepoints. Total lipids in each timepoint were extracted, resolved by TLC, and bands revealed by phosphor imaging. (E) The relative amounts of radioactive PS, PE, and PC were calculated for IM-Psd1 and Inv-Psd1 as a % of the sum of the combined radioactive signals at each timepoint (mean ± SD for *n* = 4 biological replicates from 2 clones/genotype). Significant differences (1 symbol, *P* ≤ 0.05; 2 symbols, *P* ≤ 0.01; 4 symbols, *P* ≤ 0.0001) between IM-Psd1 and Inv-Psd1 at each timepoint were calculated by unpaired student t tests.

### Robust flux through Inv-Psd1 independent of Ups2 or Mic60

To formally test the MICOS-organized *in trans* model, we took two independent KI clones of both IM-Psd1 and Inv-Psd1 KI and generated daughters lacking Mic60 or Ups2 individually or in combination. The steady state abundance of both subunits of IM-Psd1 and Inv-Psd1 was unchanged in the absence of Ups2, Mic60, or both (Figure 6A). Growth of IM-Psd1 and Inv-Psd1 yeast, with or without Ups2 and/or Mic60, was assessed on SD and SDLac plates devoid or replete with ethanolamine (Figures 6B-D). Focusing first on the parental strains containing both Ups2 and Mic60, Inv-Psd1 yeast had a slight but significant growth defect on SD and SDLac compared to IM-Psd1 (Figure 6C), suggesting there is a slight penalty for making PE on the matrix side of the IM. The absence of Mic60 resulted in a growth defect in both IM-Psd1 and Inv-Psd1 strains, agnostic of metabolic-demand (Figure 6D). Different from the overexpression model (Figure 2C, D), IM-Psd1 knock-in yeast lacking Ups2 displayed a modest growth defect on both SD and SDLac. In contrast, growth of Inv-Psd1 yeast was unaffected by the absence of Ups2 even in conditions requiring cellular energy be generated through oxidative phosphorylation (SDLac). This is notable because the *in trans* mode of activity is not available to Inv-Psd1 and the relative importance of Ups2/Mdm35-mediated transport of PS into the IM is thought to be greatest under respiratory conditions [24]. As expected (Figure 2C, D), the additional loss of Ups2 with Mic60 partially improved growth of IM-Psd1 yeast, but surprisingly, completely restored full growth capacity of Inv-Psd1 yeast. Intriguingly, ethanolamine improved growth of *mic60*Δ IM-Psd1, *mic60*Δ*ups2*Δ IM-Psd1, and *mic60*Δ Inv-Psd1 yeast on SDLac, suggesting this respiratory growth phenotype can be slightly improved with ER-derived PE.

**Figure 6.**
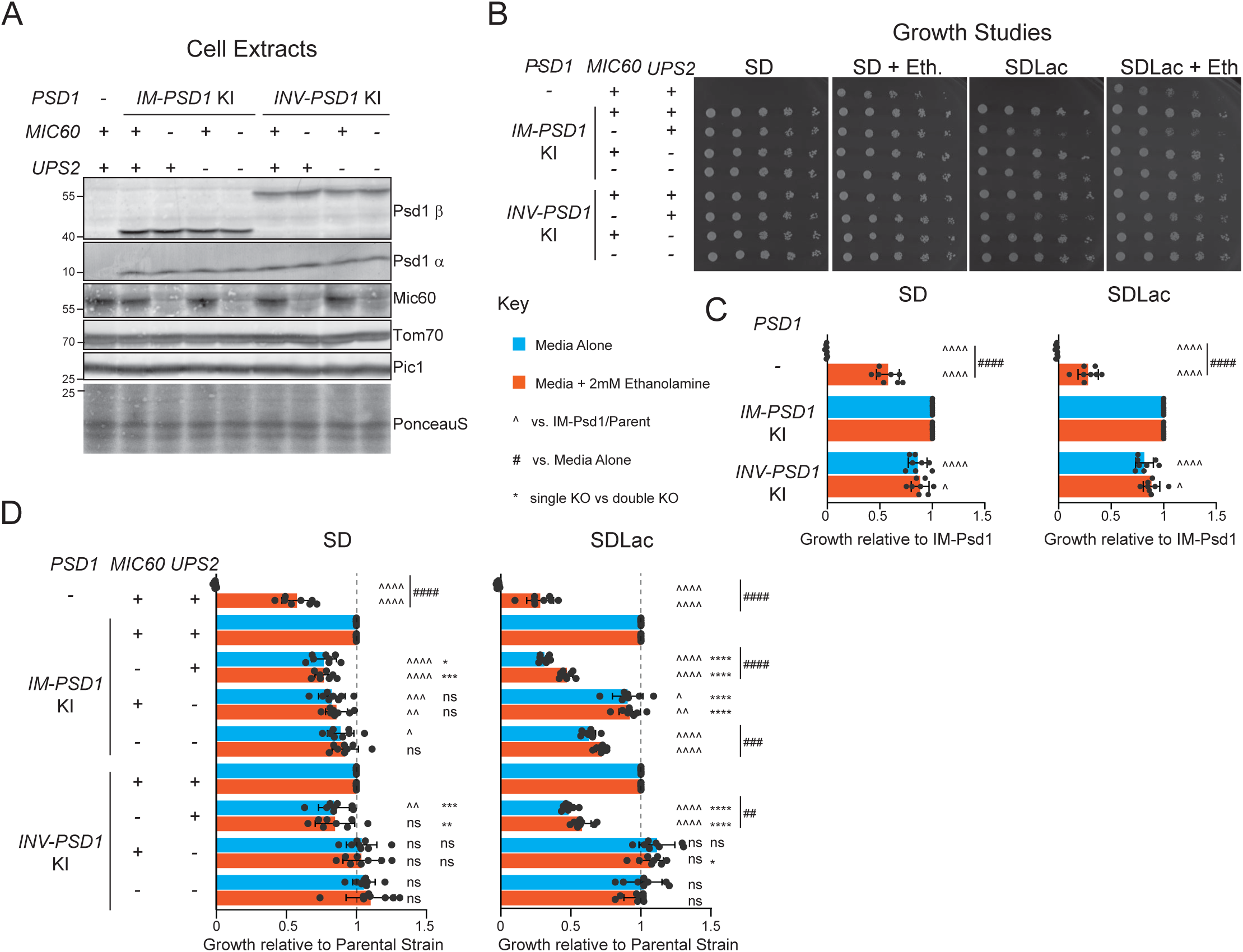
The absence of Ups2 does not impair Inv-Psd1 growth. (A) Psd1 β and α subunits were detected by immunoblot in yeast cell extracts from IM-Psd1 and Inv-Psd1 yeast of the indicated genotypes. Pic1 and Ponceau S served as loading controls. (B) The designated strains were spotted onto SD and SDLac plates, each with or without 2mM ethanolamine, and grown for 2 (SD) or 3 (SDLac) days at 30°C. (C) Growth of *psd2*Δ*psd1*Δ and Inv-Psd1 relative to IM-Psd1 (mean ± SD for *n* = 8 biological replicates). Statistical differences (ns, *P* > 0.05; 1 symbol, *P* ≤ 0.05; 2 symbols, *P* ≤ 0.01; 3 symbols, *P* ≤ 0.001; 4 symbols, *P* ≤ 0.0001) compared to IM-Psd1 were determined by one-way ANOVA with Tukey’s multiple comparisons test; differences as a function of ethanolamine were calculated by unpaired student t tests. (D) Growth of each IM-Psd1 and Inv-Psd1 daughter relative to their Ups2- and Mic60-proficient parent was determined (mean ± SD for *n* = 8 biological replicates). Statistical differences compared to the parent (^) or single versus double Ups2/Mic60 knockout (KO; *) were determined by one-way ANOVA with Sidak’s pairwise comparisons; differences as a function of ethanolamine were calculated by unpaired student t tests.

Finally, we sought to directly determine how the loss of Mic60 and/or Ups2 impacts PE metabolism in the context of either IM-Psd1 or Inv-Psd1 (Figure 7A). We first determined the steady state mitochondrial phospholipid profiles of the IM-Psd1 and Inv-Psd1 panels following overnight growth in either SD or SDLac (Figure 7B-K). Focusing first on the parental strains containing Ups2 and Mic60, compared to IM-Psd1, PS levels were increased in Inv-Psd1 yeast grown in SD while PE levels were slightly decreased in SDLac, the latter of which resulted in an elevated PC:PE ratio (Figure 7E, H, K). In both IM-Psd1 and Inv-Psd1 yeast, PE amounts were similarly decreased in the absence of Ups2 alone or in combination with Mic60, regardless of metabolic demand. These alterations in PE levels shifted the PC:PE ratios in most IM-Psd1 and Inv-Psd1 daughters compared to their Mic60- and Ups2-proficient parental counterparts (Figure 7G, K). Additional changes noted were dependent on metabolic conditions. For example, when fueled by glycolysis (SD), the absence of Mic60 resulted in a slight increase in PE in both IM-Psd1 and Inv-Psd1 yeast, whereas in respiratory conditions (SDLac), PE levels decreased (Figure 7D, H). Taken together, these results demonstrate that Ups2/Mdm35 play an important role in setting the steady state level of mitochondrial PE, regardless of the IM topology of Psd1 or the yeast metabolic state. Further, they establish that unlike Ups2/Mdm35, Mic60 and by extension MICOS, is not a major determinant of the steady state mitochondrial phospholipid profile of IM-Psd1, as seen previously [23], or Inv-Psd1 yeast.

**Figure 7.**
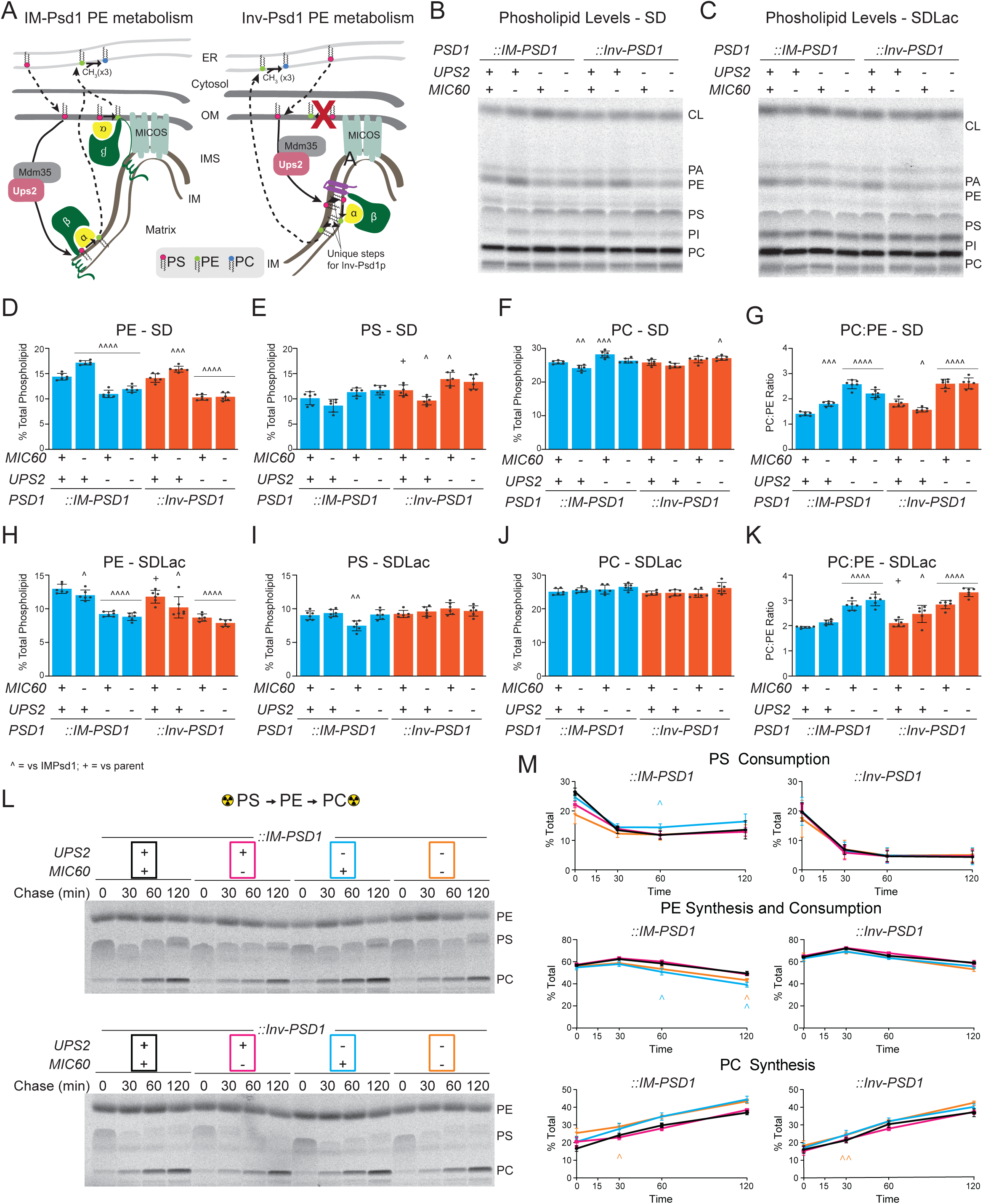
Flux through Inv-Psd1 in combined absence of Ups2 and Mic60 is unperturbed. (A) Cartoon outlining IM-Psd1 and Inv-Psd1 PE metabolism highlighting the proposed roles for Ups2/Mdm35 and Mic60-containing MICOS and the inability of Inv-Psd1 to engage the latter. Mitochondrial phospholipids were labeled overnight with ^14^C-acetate in indicated yeast grown in (B) SD or (C) SDLac medium, extracted and resolved by TLC. Quantitation of mitochondrial PE (D, H), PS (E, I), PC (F, J) levels, and the PC:PE ratio (G, K) (mean ± SD for *n* = 6 biological replicates from 2 clones/genotype). Significant differences compared to the respective Ups2- and Mic60-proficient parent (^) were determined by one-way ANOVA with Tukey’s multiple comparisons; differences between Ups2 and Mic60 containing IM-Psd1 and Inv-Psd1 (+) were determined by unpaired student t test. (L) IM-Psd1 and Inv-Psd1 yeast of indicated genotype were pulse-labeled with ^14^C(U)L-serine for 15min, washed, and then chased in SD medium with nonradioactive serine shaking at 30°C before removing aliquots at the indicated timepoints. Total lipids in each timepoint were extracted, resolved by TLC, and bands revealed by phosphor imaging. (M) The relative amounts of radioactive PS, PE, and PC were calculated as a % of the sum of the combined radioactive signals at each timepoint (mean ± SD for *n* = 4 biological replicates from 2 clones/genotype). Significant differences (1 symbol, *P* ≤ 0.05; 2 symbols) compared to the respective Ups2- and Mic60-proficient IM-Psd1 or Inv-Psd1 parent (+) were calculated by one-way ANOVA with Tukey’s multiple comparisons test.

We then measured flux through IM-Psd1 and Inv-Psd1 as a function of the presence or absence of Mic60 and/or Ups2 (Figure 7L). To minimize the relative contribution of Ups2/Mdm35, these assays were performed under glycolytic conditions (SD media). Consistent with prior work [23, 24] and the glycolytic metabolic state, flux from radiolabeled PS to PC was not impaired by loss of Ups2 in either IM-Psd1 or Inv-Psd1 yeast (Figure 7M). Flux through IM-Psd1 was also insensitive to the absence of Mic60 alone, in contrast to prior work [23], or in combination with Ups2. Surprisingly, flux through Inv-Psd1 remained intact even in the combined absence of Ups2 and Mic60. These cumulative results demonstrate that the major conduit(s) for PS, and likely other phospholipids, across the IMS has yet to be identified. This in turn suggests a new model of how Ups2/Mdm35 help establish the steady state level of PE in mitochondria via their kinetically slow, vectorial delivery of PS from the OM to the IM that offsets a comparatively fast and major bidirectional route of phospholipid transport.

## Discussion

Since it was first suggested nearly a decade ago, the proposed MICOS-organized *in trans* model of Psd1 function had sat unchallenged until now [23]. In the present study, we implemented a two-pronged rewiring-based strategy to 1) confirm that Ups2 and Mic60, the latter acting as a proxy for MICOS, function within the IMS in the context of mitochondrial PE metabolism, as expected, and 2) formally test the *in trans* model using a topologically inverted Psd1 chimera that is physically separated from the IMS-facing leaflet of the OM. Our results with OM-Psd1 demonstrate that it circumvents the requirement for either Ups2/Mdm35 or Mic60. This was the expected result and is consistent with the conclusion that with respect to mitochondrial PE metabolism, the roles performed by Ups2/Mdm35 and MICOS, if any, occur underneath the OM.

Our results testing the *in trans* model using Inv-Psd1 are much more profound and provocative. First, flux through Inv-Psd1was intact but differed from than IM-Psd1 up until the generation of PC in the ER from radiolabeled PE. This demonstrates that PS can flip to the matrix side of the IM to gain access to Inv-Psd1, and that PE, based on its subsequent trimethylation to PC, can move in the opposite direction. To our knowledge, neither trafficking step had been demonstrated previously, although a similar chimeric strategy showed that PS made in the matrix-facing IM leaflet can flip to the IMS-side [48]. Based on the observed accumulation of PE for Inv-Psd1 in the ^14^C-serine pulse-chase paradigm, we speculate that the transbilayer movement of PS between IM leaflets is faster and/or more robust than that of PE. Second, there was a slight growth penalty tied to making PE on the matrix side of the IM that did not correlate with any significant differences in the steady state phospholipid profile. The basis for this growth defect is unclear but could reflect a change in the transbilayer distribution of PE in Inv-Psd1 yeast that the methods used here are unable to capture. Third, the absence of Mic60, used here as a proxy for the loss of MICOS, resulted in a growth defect for IM-Psd1 and Inv-Psd1 yeast. In respiratory conditions (SDLac media), this *mic60*Δ-associated growth defect was accompanied by a slight but significant decrease in PE amounts and was partially ameliorated by the presence of ethanolamine. This could possibly indicate that extramitochondrial PE can normalize PE levels in the absence of Mic60 and improve growth when the carbon source provided requires mitochondrial energy production. Under glycolytic conditions however, IM-Psd1 and Inv-Psd1 yeast lacking Mic60 contained higher PE levels and their growth defects were unresponsive to ethanolamine, suggesting that either the mechanistic basis for this growth defect varies based on metabolic state or is not directly tied to phospholipid metabolism. Fourth, the additional loss of Ups2 with Mic60 partially or fully restored growth of IM-Psd1 and Inv-Psd1, respectively, and reduced the steady state amounts of PE to those found in cells lacking Ups2 alone. These results are consistent with the proposed model that additional loss of Ups2 in yeast lacking MICOS improves growth by preventing the accumulation of toxic levels of PE in the IM [23]. However, the ability of ethanolamine to improve growth of Mic60-deficient IM-Psd1 and Inv-Psd1 yeast, particularly the *mic60*Δ*ups2*Δ IM-Psd1 strain, on SDLac suggests that the basis for this growth defect and how it interfaces with PE metabolism has more mechanistic layers than this low-PE rescue model captures. Fifth, flux through IM-Psd1 under glycolytic conditions proceeds unimpeded in the combined absence of Ups2 and Mic60, the only factors directly implicated in enabling Psd1-mediated PE production in yeast. The lack of major impairment in conversion of PS to PE by IM-Psd1 when Ups2 and Mic60 are absent formally demonstrates that an(other) more significant player involved in providing Psd1 access to its substrate has yet to be identified. Sixth, flux through Inv-Psd1 in glycolytic conditions similarly proceeds unimpeded in the combined absence of Ups2 and Mic60. Given that Inv-Psd1 is physically unable to engage PS via an *in trans* mechanism, this indicates that if MICOS participates directly in mitochondrial PE metabolism, it does so via a distinct and decidedly minor mechanism. Additionally, the persistence of PE metabolism through Inv-Psd1, even without Ups2 and Mic60, further underscores the existence of a major and presently unknown conduit for PS, and likely other phospholipids, across the IMS.

The lack of an effect on PE metabolic flux when Ups2 is absent seems, on the surface, to contradict the clear and metabolic state-independent drop in steady state PE amounts in both IM-Psd1 and Inv-Psd1 yeast. The existence of an unknown and kinetically more robust pathway for PS to traverse the aqueous IMS reconciles this paradox. Based on our prior [49] and current results, we propose the following refined framework for mitochondrial PE metabolism in yeast. There is at least one proportionately major route for PS across the IMS yet to be molecularly identified. We postulate that this major conduit is likely nonspecific in terms of its cargo and bidirectional, allowing PS to move in both directions through this medium. In this context, vectorial delivery of PS across the IMS is conferred by the combined actions of 1) Psd1-mediated consumption of PS in the IM, which helps maintain a PS gradient from the ER to the IM; and 2) kinetically slow Ups2/Mdm35-mediated delivery of one PS molecule per Ups2/Mdm35 dimer from the OM to the IM [25, 26]. The contribution of Ups2/Mdm35 to steady state PE levels is clear in the nearly 35-50% decrease in PE observed when its activity is lost. The lack of clear effect on flux in the context of a 135min pulse-chase protocol likely reflects that directionality of PS delivery by Ups2/Mdm35 is regulated through several biochemical mechanisms. This is supported in several ways. In addition to providing PS scramblase activity [22], Por1 also serves as a scaffold that facilitates the productive coupling of Mdm35 with Ups2 [63]. Once formed, each Ups2/Mdm35 dimer can accommodate a single PS molecule and transport it across the aqueous IMS between the two mitochondrial membranes [23–26]. Directionality is further provided by its preferred association with CL-containing membranes which also favors the delivery and release of PS in the IM [23]. The proposed mechanism of CL-facilitated PS release also involves the dissociation of Ups2 from Mdm35 [23], the former of which is then susceptible to degradation by the IM-resident protease, Yme1 [27]. Thus, based on what is known, each individual Ups2 protein may be limited to only a single OM to IM PS transport event without the chance for additional trips. And yet, at steady state, through this seemingly incremental delivery of PS into the IM, Ups2/Mdm35 play a significant role in establishing the membrane diversity of the IM. The molecular identification of the currently unknown entity responsible for the kinetically dominant movement of PS across the IMS is a major goal of future research that once achieved, will allow our new model of mitochondrial PE metabolism to be formally put to the test.

## Materials and Methods

### Yeast strains and growth conditions

The lineage of all yeast strains employed in this study was derived from GA74-1A (*MAT*a, *his3-11,15*, *leu2*, *ura3*, *trp1*, *ade8* [rho^+^, mit^+^]). The *psd2*Δ (*MAT*a, *psd2::HISMX6*, *leu2*, *ura3*, *trp1*, *ade8* [rho^+^, mit^+^]) and *psd2*Δ*psd1*Δ (*MAT*a, *psd2::HISMX6*, *leu2*, *ura3*, *psd1::TRP1*, *ade8* [rho^+^, mit^+^]) yeast strains were previously described [56]. The pRS305 plasmids containing wild type Psd1 (referred to as IM-Psd1 in this study), autocatalytic mutant (461-LGS/AAA-463) Psd1, or OM-localized Psd1, each with a COOH-terminal 3XFLAG tag, and the *psd2*Δ*psd1*Δ-based strains transformed with them, have been previously described [49, 56]. To topologically invert Psd1 to face the mitochondrial matrix, the first 105 amino acids containing the mitochondrial targeting sequence and transmembrane domain of Psd1 were replaced by residues 1-128 of Bcs1, a single pass IM protein with its C-terminus in the matrix [48]. Inv-Psd1 containing a C-terminal 3XFLAG tag (DYKDHDGDYKDHDIDYKDDDDK) and IM-Psd1 and Inv-Psd1 harboring C-terminal SpyTag2 (VPTIVMVDAYKRYK) were generated by overlap extension PCR and subcloned into pRS305. Autocatalytic-dead Inv-Psd1 was generated by subcloning Inv-Psd1 into pRS305Psd1^LGS/AAA^3XFLAG. All pRS305-based Psd1 constructs were linearized with AflII and integrated into the LEU2 locus of the *psd1*Δ*psd2*Δ yeast strain. Clones were selected on synthetic dropout medium (0.17% (w/v) yeast nitrogen base (US Biological Y2035), 0.5% (w/v) ammonium sulfate, 0.2% (w/v) dropout mixture synthetic-leu (US Biological D9526), 2% (w/v) dextrose, 2% (w/v) agar) -leucine and verified by immunoblot. To adapt the SpyCatcher/SpyTag system [58] to probe the IM topology of Inv-Psd1, SpyCatcher002 and SpyCatcher002 EQ, a non-reactive control unable to form an isopeptide bond upon engaging SpyTag002, were initially subcloned from pDEST14-SpyCatcher002 (a gift from Mark Howarth (Addgene plasmid # 102827; http://n2t.net/addgene:102827; RRID:Addgene_102827) and pDEST14-SpyCatcher002 EQ (a gift from Mark Howarth (Addgene plasmid # 102830; http://n2t.net/addgene:102830; RRID:Addgene_102830) [57] into pFA6a-KanMX6 (a gift from Jurg Bahler & John Pringle (Addgene plasmid # 39296; http://n2t.net/addgene:39296; RRID:Addgene_39296) [64]. Using overlap extension PCR, C-terminal His_8_ tags were added to SpyCatcher002 and SpyCatcher002 EQ (amplified using pFA6aSpyCatcher002-KanMX6 or pFA6aSpyCatcher002 EQ-KanMX6 as templates) which were in turn each placed downstream of the matrix-destined mitochondrial targeting signal from subunit 9 of complex V from *Neurospora crassa* (amplified using pRS313-Su9-GFP [65]; a kind gift of Hiromi Sesaki, JHMI). Su9SpyCatcher002-His and SpyCatcher002 EQ-His were placed under the control of the 5’ and 3’ UTRs of *PSD1* (amplified using pRS305Psd1-3XFLAG as template) and subcloned into pRS316. All newly generated plasmids were sequence verified. pRS316-based yeast transformants were selected and maintained on synthetic dropout medium (0.17% (w/v) yeast nitrogen base, 0.5% (w/v) ammonium sulfate, 0.2% (w/v) dropout mixture synthetic-uracil (US Biological D9536), 2% (w/v) dextrose) -Uracil; solid formulations included 2% (w/v) agar.

To generate IM-Psd1 and Inv-Psd1 knock-in (KI) strains, targeting constructs were produced by PCR amplification using primers that annealed in the 5’ or 3’ UTRs of *PSD1* and pRS305IMPsd1 or pRS305Inv-Psd1, each with 3XFLAG tags, as templates. Following transformation of *psd2*Δ*psd1*Δ yeast with 3 µg of each targeting construct, potential KI strains were identified by counter-selecting against the presence of *TRP1* on 5-Fluoroanthranilic acid (5-FAA; 5% (w/v) glucose, 0.67% (w/v) yeast nitrogen base with 0.5% (w/v) ammonium sulfate (RPI Y20040), 0.0285% (w/v) dropout mix complete (US Biological D9516), 0.08575 % (w/v) dropout mix synthetic-tryptophan (US Biological D9531), 0.1g/mL 2-Amino-5-fluorobenzoic acid (5-FAA; added from a 10% (w/v) ethanolic stock post-autoclaving; Sigma 367982), 2% (w/v) agar) plates and verified by immunoblot.

Deletion of the *UPS2* and *DPL1* genes, and for the OM-Psd1 strains, the *MIC60* gene, was achieved through genetic modification via homology-integrated clustered regulatory interspaced short palindromic repeats (CRISPR)-Cas (HI-CRISPR) [66]. CRISPR-Cas9 gene blocks were designed targeting *DPL1*, *UPS2*, or *MIC60* and assembled into the pCRCT plasmid (pCRCT was a gift from Huimin Zhao (Addgene plasmid #60621; http://n2t.net/addgene:60621; RRID:Addgene_60621), as described in [13, 49]. The gene blocks included the 20bp CRISPR-Cas9 target and the homology repair template with homology arms spanning 50bp on each side to sum to 100bp flanking the Cas9 recognition sequence and were ordered as Geneblocks (Integrated DNA Technologies). Each knockout was achieved by a homology repair template that contained an 8bp deletion near the N-terminus to both induce a frameshift mutation that results in a premature stop codon and to remove the PAM sequence to prevent re-cleavage by Cas9 after the homology directed repair occurs. These 8 nucleotides were encoded by nucleotides 134-153 for the *DPL1* knockout, 140-159 for the *MIC60* knockout, and 231-250 for the *UPS2* knockout; nucleotide numbers are with respect to the open reading frame (ORF) start codon ATG sequence, defined as 1-3. The CRISPR-Cas9 system targeted the protospacer adjacent motif (PAM) sequence encoded by nucleotides 154-156 on the + strand of the ORF for Dpl1, 239-258 on the + strand of the ORF for Mic60, and 251-253 on the + strand of the ORF for Ups2. Successful deletion of these genes following transformation and selection was verified by immunoblotting for Mic60 and PCR and subsequent sequencing of yeast genomic DNA using primers specific for *DPL1* and *UPS2*. Due to a low deletion efficiency using HI-CRISPR, *MIC60* was disrupted in the IM-Psd1 and Inv-Psd1 KI strains by replacing the entire open reading frame of *MIC60* with *TRP1*. Growth and expression of individual clones per genotype were systematically compared to each other to ensure that representative phenotypes were recorded prior to all comparisons performed across genotypes.

Other than the pRS316-based transformants (maintained on synthetic-Uracil plates), yeast were maintained on rich lactate plates (1% (w/v) yeast extract, 2% (w/v) tryptone, 0.05% (w/v) dextrose, 2% (v/v) lactic acid, 3.4mM CaCl_2_-2H_2_O, 8.5mM NaCl, 2.95mM MgCl_2_-6H_2_O, 7.35mM KH_2_PO_4_, 18.7mM NH_4_Cl, pH 5.5, 2% (w/v) agar). For growth studies, starter cultures were grown overnight in liquid rich lactate and 0.008 OD_600_ of cells was diluted with sterile ddH_2_O for Vol_final_ of 200µL followed by four 1:4 serial dilutions in sterile ddH_2_O. 3uL of cells from each dilution were spotted onto synthetic defined dextrose (SD; 0.67% (w/v) yeast nitrogen base with 0.5% (w/v) ammonium sulfate (RPI Y20040), 2% (v/v) complete amino acid mixture (added post-autoclave from a 50X filter-sterilized stock that contained 1g/L Adenine, 1g/L L-Arginine, 1g/L L-Histidine, 3g/L L-Leucine, 11.5g/L L-Lysine, 1g/L L-Methionine, 15g/L L-Threonine, 1g/L L-Tryptophan, 1g/L L-Uracil), 2% (w/v) dextrose, 2.5% (w/v) agar (Sigma 05038 or HiMedia RM301)) or synthetic defined lactate (SD-LAC; 0.67% (w/v) yeast nitrogen base with 0.5% (w/v) ammonium sulfate, 2% (v/v) complete amino acid mixture, 0.05% (w/v) dextrose, 2% (v/v) lactic acid, 3.4mM CaCl_2_-2H_2_O, 8.5mM NaCl, 2.95mM MgCl_2_-6H_2_O, 7.35mM KH_2_PO_4_, 18.7mM NH_4_Cl, pH 5.5, 2.5% (w/v) agar) agar plates, both formulations +/-2mM sterile-filtered ethanolamine hydrochloride, wrapped in parafilm and grown at 30°C. Images of yeast plates were captured face-up with the lids removed using a Chemidoc MP Imaging System (BioRad) and colorimetric acquisition mode. To quantitatively compare growth, the background subtracted volume (INT*mm) from two dilutions per genotype were averaged and growth relative to a control genotype on the same plate determined.

### Preparation of yeast cell extracts

Other than the pRS316-based transformants, all yeast were grown overnight in 2mL rich lactate; the pRS316-based transformants were grown overnight in 2mL synthetic lactate-Ura (0.67% (w/v) yeast nitrogen base with 0.5% (w/v) ammonium sulfate, 0.2% (w/v) dropout mixture synthetic-uracil, 0.05% (w/v) dextrose, 2% (v/v) lactic acid, 3.4mM CaCl_2_-2H_2_O, 8.5mM NaCl, 2.95mM MgCl_2_-6H_2_O, 7.35mM KH_2_PO_4_, 18.7mM NH_4_Cl, pH 5.5). The next day, 1 OD_600_ of cells were sedimented in 1.5mL microcentrifuge tubes by centrifugation for 10 min at 845 x *g* at either room temperature (harvested immediately) or 4°C (stored at -20°C post-aspiration). The cell-free media was aspirated and the cell pellet resuspended in 1mL of water. 150µL of a freshly made NaOH/β-mercaptoethanol solution (1mL 2M NaOH: 80µL of β-mercaptoethanol) was added to the tube which was in quick succession, mixed by inversion and placed in ice. Following a 10 min incubation on ice with regular inversions every 2 min, 75µL of 100% (w/v) tricholoroacetic acid (TCA) was added and the tube was mixed by inversion, re-placed in ice and again incubated for 10 min with regular mixing every 2 min. Following a 2 min centrifugation at 21,000 *x g* (room temperature or 4°C) to collect proteins, the supernatant was aspirated and 1mL of acetone was added to wash the protein pellet. Following another 2 min centrifugation at 21,000 *x g*, the acetone wash was decanted and the remaining pellet was briefly dried with the tube inverted on the bench before 30µL of 0.1M NaOH was added to the pellet. After each pellet had begun to dissolve at room temperature, an equal volume of 2X reducing sample buffer was added, the sample was finger-flicked and then boiled for 5 min at 95°C. Equal volumes (ranging from 6 to 8µL depending on target protein) of yeast extracts were loaded onto SDS-PAGE for immunoblotting.

### Immunoblotting

Protein samples in reducing sample buffer were resolved on homemade 10-16% or 10-18% gradient SDS-PAGE gels. Proteins were transferred post-resolution onto PVDF (Millipore Immobilon-FL, 0.45µM Catalog No. IPFL00010) with 1X Transfer Buffer (25mM Tris, 192 mM Glycine, 2.5% (v/v) methanol) at 30 Volts overnight (18hrs) at room temperature. The transfer was checked via Ponceau S staining and membrane strips were cut to probe for proteins based on their in-house verified molecular weight. As described previously [67, 68], membranes were blocked with 5% (w/v) milk (Giant, Ellicott City, MD), 0.05% (v/v) Tween-20/1XPBS for 1 hr and subsequently incubated with primary antibody while rocking for 1 hr at room temperature. Following three 10 min washes with PBST (PBS with 0.2% (v/v) Tween-20), blots were incubated with Starbright 520 or 700-conjugated secondary antibodies for 45 min, the membranes were washed thrice for 10 min with PBST, twice for 10 min with PBS and placed on paper towels to dry overnight protected from ambient light. Fluorescent blots were imaged using a BioRad ChemiDoc MP imaging system. On occasion, membranes were reprobed with a different primary antibody/secondary antibody combination. In these cases, membranes were re-wet with methanol and subjected to the entire immunoblotting regimen again, but utilizing a secondary antibody conjugated to a different Starbright dye than originally used.

### Lipid Steady States

For ::IM-Psd1 and ::OM-Psd1 analyses (Fig. 2), starter cultures grown in 2 mL of rich lactate medium at 30°C for 2-3 days, depending on the genotype, were used to inoculate 2mL YPD for final OD_600_ = 0.05 or 2mL rich lactate for final OD_600_ = 0.4. For IM-Psd1 KI and Inv-Psd1 KI analyses (Fig. 7), starter cultures grown in 3mL of rich lactate medium at 30°C for 2 days were used to inoculate 2mL SD for final OD_600_ = 0.09 (based on delayed growth, a final OD_600_ = 0.11 was used for *mic60*Δ strains) or 2 mL SD-LAC for final OD_600_ = 0.18 (final OD_600_ = 0.36 and 0.24 for *mic60*Δ and *mic60*Δ*ups2*Δ strains, respectively). Lipid labeling was initiated by sterilely adding 1µCi ^14^C-acetate (Perkin Elmer, NEC084H001MC) to each culture. After growing for 24 hrs in a water bath shaking at 240 rpm at 30°C, yeast were harvested by spinning at 3000 rpm in a clinical centrifuge for 5 min at room temp. Radioactive supernatants were aspirated, the pellets washed with 2mL sterile water, and yeast collected as before. Aspirated yeast pellets were resuspended in 0.3mL MTE buffer (0.65M Mannitol, 20mM Tris, 1mM EDTA) spiked with protease inhibitors (1mM PMSF, 10µM leupeptin, and 2µM pepstatin A) and transferred to 1.5mL microcentrifuge tube containing ∼ 0.1mL glass beads. After parafilm-sealing each, the yeast were mechanically ruptured by vortexing on high for 35 min in a cold room. Following a 376 *x g* spin at 4°C, the supernatants were transferred to new 1.5mL microcentrifuge tubes and crude mitochondria harvested by centrifugation at 13,000 x *g* for 5 minutes at 4°C. Post-aspiration, the mitochondrial-enriched pellets were resuspended by repeat pipetting with 50µL BB7.4 (0.6M Sorbitol, 20mM HEPES-KOH, pH 7.4) and ^14^C-acetate incorporation measured in 1µL by liquid scintillation counting. Phospholipids from mitochondria normalized by their ^14^C-acetate incorporation were extracted in 5mL borosilicate tubes with 1.5mL 2:1 Chloroform:Methanol by vortexing on medium-high for 30 minutes at room temperature. To improve eventual lipid migration on thin layer chromatography (TLC) plates, ∼0.5mg cold mitochondria was added to each sample prior to lipid extraction. To initiate phase separation, 0.3mL of Normal Saline (0.9% (w/v) NaCl) was added and the samples vortexed for an additional 1 min. Samples were spun in a clinical centrifuge at 1000 rpm for 5 min at room temperature and the upper aqueous phase aspirated. Organic phases were washed with 0.25mL 1:1 Methanol:H_2_O, vortexed for 30 sec, and phases separated as in the prior step. Lower organic phases were transferred to new 5mL borosilicate tube, dried down under a stream of nitrogen, and either analyzed directly by TLC or stored at -20°C until ready to resolve. Machery-Nagel SILGUR 25 TLC plates were washed with chloroform, air dried, pre-treated with 1.8% (w/v) boric acid in 100% ETOH, and then activated at 95°C for at least 30 min. Lipid samples were resuspended in 40µL chloroform and 13µL of each loaded onto TLC plates using a CAMAG Linomat 5. Plates were resolved once in equilibrated TLC tanks containing 30:35:7:35 (v/v) chloroform:ethanol:water:triethylamine, air dried, exposed to phosphoimaging K screens, and signals detected using a Molecular Dynamics Storm 820 Phosphor Imager.

### Subcellular fractionation and mitochondrial Isolation

Mitochondrial isolation and subcellular fractionation was carried out as described previously [68]. SpyTag/SpyCatcher strains were cultured in synthetic lactate-Ura whereas the IM-Psd1 and Inv-Psd1 KI strains were grown in rich lactate. Precultures grown at 30°C for 36-48hrs were used to sterilely inoculate two separate 2L flasks/strain containing 950mL synthetic lactate-Ura or rich lactate with ∼100 OD_600_’s. Inoculated final cultures were grown shaking at 220 rpms at 30°C overnight. The next day, cultures had typically reached an OD_600_ of 1.1‒1.4 and were checked for contamination. Cells were then harvested by centrifugation for 5 min at 6,000 *x g* at room temperature using 1000mL buckets. The yeast pellets were then resuspended in ∼100mL of water and transferred to pre-weighed 250mL bottles. Cell slurries were centrifuged at 2000 *x g* for 5 min at room temperature, the supernatant decanted, and the cell pellets weighed. Cell pellets were then resuspended in 50mL of freshly prepared 0.1M Tris-SO_4,_ pH 9.4 (1M Tris with pH adjusted with sulfuric acid) containing 15mM dithiothreitol, and incubated for 20 min at 30°C shaking at 220rpm. Afterward, this cell suspension was centrifuged at 2000 *x g* for 5 min at room temperature, washed with 40mL 1.2M sorbitol, 20mM KPi, pH 7.4 buffer, and yeast pellets collected again at 2000 *x g* for 5min at room temperature. After decanting the supernatant, yeast pellets were converted to spheroplasts by resuspending in 1.2M sorbitol, 20mM KPi, pH 7.4, containing 3mg of Zymolyase 20T (Nacalai Tesque, INC.) per gram of yeast, using a final volume of 2mL per gram of yeast. These suspensions were then shaken at 220rpm for ∼1hour at 30°C. Spheroplasting efficiency was monitored under a microscope by checking for yeast lysis upon dilution in 10µL water. After confirming that cells were sensitive to osmotic changes, spheroplasts were collected by centrifugation at 3500 *x g* for 5 min at 4°C. Moving forward, cells were kept on ice or at 4°C. Cell pellets were washed twice with 1.2M sorbitol, 20mm KPi, pH 7.4 buffer chilled at 4°C and collected by centrifugation at 3500 *x g* for 5 min at 4°C. Spheroplasts were resuspended in 50mL of 0.6M sorbitol, 20mM KOH-MES, pH 6.0 (BB6.0 buffer) containing 1mM phenylmethylsulfonyl fluoride (PMSF) and decanted into a tight-fitting (type A) glass dounce homogenizer kept on ice where they were homogenized using 15 strokes. For cell fractionation studies, 200µL of this homogenate was collected and placed on ice to quantitate and analyze as the starting material (SM). The combined homogenates were centrifuged for 5 min at 1700 *x g* at 4°C, and the supernatants collected while the residual pellets were re-homogenized in BB6.0 + 1mM PMSF by 15 strokes in the same glass dounce to extract more membrane-bound organelles. This second round of homogenate was centrifuged at 1700 *x g* at 4°C and the supernatants from both homogenizations were combined and centrifuged for 10 min at 13,500 *x g* and 4°C. These subsequent pellets were enriched for mitochondria. The supernatants (S13) were either further fractionated (described below) or discarded. The mitochondria-enriched pellets were washed with 35mL of BB6.0 buffer lacking PMSF. Mitochondria were resuspended 2 times using a pre-chilled Teflon dounce and following a 1700 *x g* 5 min spin at 4°C, the mitochondria-containing supernatant was transferred to fresh 50mL tube; the pellets were discarded. Supernatants were centrifuged at 13,500 *x g* for 10min at 4°C. This time, the supernatant was aspirated, and the mitochondrial pellet was washed once with ∼30mL of BB7.4 buffer. Mitochondria were again resuspended 2 times using the pre-chilled Teflon dounce, transferred to a clean 50mL tube, and harvested at 13,500 *x g* for 10min at 4°C. The final crude mitochondrial pellet (P13) was concentrated by aspirating the supernatant and the mitochondrial pellets were resuspended in residual BB7.4. Protein concentration was determined using the Pierce BCA Protein Assay Kit (ThermoFisher Scientific, Catalog No. 23225). Aliquots of mitochondria (1 mg at ∼25mg/mL) were snap frozen in liquid nitrogen and stored at -80°C.

Additional subcellular fractions were collected starting with 35mL of the S13 supernatant, which was transferred to a 50mL tube and centrifuged at 21,500 *x g* for 15 min at 4°C. The resulting supernatant was then transferred to a fresh 50mL tube and spun at 40,000 *x g* for 30min at 4°C. The generated cytosol and light-membrane containing supernatant (S40) was transferred to a 50mL falcon tube. The ER-enriched 40,000-x *g* pellet (P40) was resuspended in residual buffer and transferred to a 1.5mL microcentrifuge tube. The protein concentration of each collected fraction (SM, P13, P40, and S40) was determined with the BCA assay. Aliquots of cell fractionation samples were snap frozen in liquid nitrogen and stored at -80°C.

### Protease protection assay

An established protease accessibility assay [69] was used to determine the IM topologies of IM-Psd1 and Inv-Psd1. This assay determines the protease-sensitivity of proteins as a function of the integrity of mitochondrial membranes: intact mitochondria were maintained in an iso-osmotic buffer, OM-disrupted mitoplasts were incubated in hypo-osmotic conditions which causes the IM to unfurl and physically rupture the OM, or mitochondria were solubilized with detergent. For intact mitochondria, two microcentrifuge tubes containing 150 µg of mitochondria were set aside on ice. For OM-disrupted/detergent-solubilized samples, 600 µg of mitochondria was centrifuged at 8,000 *x g* for 5 min at 4°C. Post-aspiration, the mitochondrial pellet was resuspended in 200µL of BB7.4 and 50 µL was distributed into four tubes each. To rupture the OM, 19x volumes of 20mM K^+^HEPES, pH 7.4, with or without 100µg/mL of Proteinase K, was added to the mixture containing mitochondria. For detergent solubilization, 19x volumes of 20mM K^+^HEPES, pH 7.4 with 0.5% (w/v) deoxycholate, with or without 100µg/mL Proteinase K, was added. In parallel, the intact mitochondrial samples were resuspended in 1mL BB7.4 buffer with or without 100µg/mL Proteinase K. After mixing via vortex on low-medium for 15sec, the samples were incubated on ice for 30 min. Once done, PMSF was added (5mM final) to each sample to inhibit Proteinase K, and then samples were centrifuged at 21,000 *x g* for 10 min at 4°C. For intact mitochondria and OM-ruptured mitoplasts, the supernatant was aspirated and Proteinase K was completely inactivated as follows: pellets were resuspended in 180µL of BB7.4 containing 1mM PMSF and transferred to fresh tubes containing 20µL of 100% (w/v) TCA which were heated at 60°C for 5 min and then placed on ice for at least 5 min. For detergent-solubilized mitochondria, the supernatants were transferred to tubes containing 0.2mL of 100% (w/v) TCA, heated at 60°C for 5 min and then incubated on ice for 1 hr. Following these steps, all samples were centrifuged at 21,000 *x g* for 10 min at 4°C, the supernatants aspirated, and pellets washed with 0.5mL of cold acetone. The acetone was decanted following a 10 min 21,000 *x g* centrifugation at 4°C and the pellets incubated with 30µL of 0.1M NaOH for 30min at room temperature. An equal volume of 2X reducing sample buffer was added to each sample. Samples were then boiled for 5 min at 95°C. 20µg of each sample was loaded and evaluated by SDS-PAGE and immunoblotting.

### NBD-PS decarboxylation assay

The Psd1 enzyme activity was measured through NBD-PS fluorescence, as previously reported [22, 24], but with slight modifications. Briefly, mitochondria (400µg) were centrifuged at 8,000 x *g* for 5 min at 4°C, and the supernatant aspirated. The pellets were resuspended in 10mM MES pH 6.0, 0.6M Sorbitol and 1.55mM TX-100 by pipetting for a final concentration of 2.5 mg/ml, and vortexed briefly on low followed by incubation on ice for 5 min. To initiate the Psd1 enzymatic reaction, mitochondria were further diluted to 0.5 mg/mL with assay buffer (10mM MES pH 6.0, 0.6M Sorbitol, 12.5mM EDTA, 2μM 16:0-C12 NBD PS; Avanti Polar Lipids, Inc., 81093C). Samples were incubated at 30°C, and at the indicated timepoints, the reaction was terminated by transferring 100µL aliquots to microfuge tubes containing 800µL 2:1 chloroform:methanol, vortexing briefly on high and incubating on ice until all the timepoints were collected. To initiate phase separation, 200µL 0.9% (w/v) NaCl was added and the samples vortexed at RT for 5 min, followed by centrifugation at 2180 x *g* for 5 min. After aspirating the upper aqueous phase, samples were dried using a N-EVAP Nitrogen Evaporator (Organomation). Samples were resuspended in 13µL chloroform immediately prior to loading onto a TLC plate (TLC silica gel 60; MilliporeSigma Supelco 1057210001) pretreated with 1.8% (w/v) boric acid/100% ethanol and activated at 95°C for at least 30 min. TLC plates were resolved with chloroform:ethanol:water:triethylamine (30:35:7:35, by volume). NBD signal was detected using Amersham Typhoon Imager with Cy2 emission filter and quantified using ImageLab (Bio-Rad). The enzymatic conversion was assessed using the following equation:

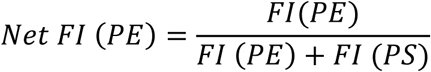

Where FI= Fluorescence Intensity.

### 14C-Serine Pulse-Chase PS-PE-PC Conversion Assay

Starter cultures were grown in 20 mL rich lactate shaking at 220rpm at 30°C for 1-2 days. On the day of the experiment, 16 OD_600_’s of each culture was transferred to sterile 17×100mm round bottom tubes, centrifuged in a clinical centrifuge at 1,690 *x g* for 5 min, the supernatant aspirated, and the pellet resuspended in 1mL SD-inositol (0.67% (w/v) Yeast Nitrogen Base with Ammonium Sulfate without Inositol (MP Biomedicals 4027412), 2% (w/v) dextrose (Sigma G8270), and post-autoclave, 2% (v/v) complete amino acid mixture). The resulting yeast slurry was added to a flask containing 19mL SD-inositol which was incubated at 30°C shaking at 220 rpm until an OD_600_ of 0.8-1.0 was reached (typically 2-4hrs). 12 OD_600_’s of each SD-inositol acclimated culture was transferred to sterile 17×100mm round bottom tubes and yeast pellets collected by centrifugation as before. Resulting supernatants were aspirated and the pellets were resuspended with 4mL SD-inositol spiked with 2.5µCi/mL ^14^C(U)L-serine (Moravek Inc., MC265 250UI) and incubated for 15min at 30°C, 240 rpm. To delay further membrane-mediated metabolism of ^14^C(U)L-serine, 4mL ice-cold SD-inositol + 10mg/mL cold L-serine was added to tubes and the yeast pellets collected by centrifugation at 1,690 *x g* at 4°C for 2 min. Supernatants were aspirated and pellets were washed with 4mL ice-cold SD-inositol + 10mg/mL cold L-serine and collected as in the prior step. Recovered ^14^C(U)L-serine pulse-labeled pellets were resuspended with 4mL pre-warmed SD-inositol + 10mg/mL cold L-serine and incubated in a shaking water bath (240 rpm) at 30°C for two hours. At the designated chase timepoints, starting at t=0, 0.97mL aliquots were transferred to fresh microcentrifuge tubes placed on ice, yeast pellets were collected by centrifugation at 21,000 *x g* at 4°C for 2 min, the supernatants aspirated, and tubes stored at -20°C until all timepoints had been collected and were ready to harvest. When ready to harvest, pellets were washed with 1mL ice-cold water and sedimented with a 2.5 min, 845 *x g* spin at 4°C. Aspirated pellets were resuspended in 0.3mL MTE buffer with protease inhibitors and mechanically disrupted with glass beads as for lipid steady state analyses. Following a 376 *x g* spin at 4°C, the supernatants were transferred to new 1.5 mL microcentrifuge tubes and ^14^C(U)L-serine incorporation measured in 1 µL by liquid scintillation counting. Phospholipids from mitochondria normalized by their ^14^C(U)L-serine incorporation were again combined with 0.5 mg cold carrier mitochondria and extracted in 5 mL borosilicate tubes with 1.5 mL 2:1 Chloroform:Methanol by vortexing on medium-high for 30 minutes at room temperature. Phase separation, organic phase washing, and organic phase collection and desiccation was done as for lipid steady states. Sample resolution by TLC was done using same system as for lipid steady states except that 100% of each timepoint was resolved and analyzed. The relative abundance of PS, PE, and PC at each timepoint was calculated as a % of the sum of the combined radioactive signals from PS, PE, and PC. Since the decarboxylation of PS to PE involves the loss of 1 of 3 labeled carbons in ^14^C(U)L-serine, the volumes of PE and PC were multiplied by 1.5.

### Antibodies

The custom produced antibodies against yeast Mic60 were generated by Pacific Immunology (Ramona, CA) using affinity-purified His_6_Mic60 as an antigen. In brief, the predicted mature Mic60 open reading frame starting at Glu-82 was cloned downstream of the His_6_ tag encoded in the pET28a plasmid (Novogene) and transformed in BL21(RIL) *Escherichia coli*. A 1L 2X YT culture containing 20µg/mL kanamycin was induced with 0.5 mM IPTG at 25°C for 4 h and the bacterial pellet collected at 3020 × *g* for 10 min, washed with 0.9% (w/v) NaCl, and stored at −20°C until purification was performed. The bacterial pellet was resuspended in 40mL lysis buffer (50mM NaH_2_PO_4_, 300mM NaCl, 10mM Imidazole, 0.1mM EDTA, pH 8.0) containing 1mg/mL lysozyme and incubated with rocking for 30 min at 4°C. The resulting bacterial suspension was obliterated using an Avestin Homogenizer and the resulting lysate centrifuged at 10,000 × *g* at 4°C for 20 min. The His_6_Mic60-containing pellet was solubilized with 5mL freshly made Inclusion Body solubilization buffer (1.67%(w/v) Sarkosyl, 0.1mM EDTA, 10 mM DTT, 10 mM Tris-Cl pH 7.4, 0.05% (w/v) PEG3350) by vortexing on high and then incubated on ice for 20 min. 10 mL of 10mM Tris-Cl pH7.4 was added to the suspension, which was then centrifuged at 12,000 × *g* at 4°C for 10 min. His_6_Mic60 was then purified from this suspension by incubating with 1.5 ml Ni-NTA in 15mL falcon tube rotating for 2hr at 4°C. The Ni-NTA mixture was loaded onto a chromatography column and the non-binding flow through collected. Following four 10mL column-volume washes with 1) 0.1% (w/v) Sarkosyl, 50mM NaH_2_PO_4_, 300mM NaCl, 20 mM Imidazole, 10% (v/v) glycerol, 20mM β-ME, pH 8.0; 2) 0.15% (w/v) Sarkosyl, 50mM NaH_2_PO_4_, 600mM NaCl, 30mM Imidazole, 20% (v/v) glycerol, 20mM β-ME, pH 8.0; 3) 0.2% (w/v) Sarkosyl, 50mM NaH_2_PO_4_, 600mM NaCl, 40mM Imidazole, 30% (v/v) glycerol, 20mM β-ME, pH 8.0; and 4) 0.1% (w/v) Sarkosyl, 50mM NaH_2_PO_4_, 300mM NaCl, 20 mM Imidazole, 10% (v/v) glycerol, 20mM β-ME, pH 8.0, bound material was recovered with elution buffer (250mM imidazole, 0.1% (w/v) Sarkosyl, 50mM NaH_2_PO_4_, 300mM NaCl, and 10% (v/v) glycerol, pH 8.0; 6 sequential 0.5mL elutions). Protein-containing fractions were identified using the Bradford Assay (Bio-Rad) and combined, PBS dialyzed, and quantified using a BSA standard curve prior to antibody generation.

Other in-house antibodies generated in either in our laboratory or the laboratory of C. Koehler (UCLA) and used in this study include rabbit anti-yeast Psd1 (Psd1β-specific; 4077.5 and 4078.5; 1:1000; [29]), rabbit anti-yeast Pic1 (3676.3; 1:10,0000; [70]), rabbit anti-yeast Mic60 (20450.F; 1:5000; this study), rabbit anti-yeast Por1 (425; 1:10,000; [71]), rabbit anti-yeast Tom70 (7306.F; 1:10,0000; [72]), rabbit anti-yeast Taz1 (4248.F; 1:1000; [68]), rabbit anti-yeast Abf2 (5477.2; 1:8000; [49]), and rabbit anti-yeast Hsp70 (SH1-T; 1:10,000; [68]). Additional antibodies employed were rabbit anti-yeast Mic60 (αFcj1; 1:1000; [53]), mouse anti-FLAG (clone M2; 1:5000; Sigma F3165;), mouse anti-His (1B7G5; 1:3000-5000; ProteinTech 66005-1), mouse anti-Dpm1 (5C5A7; 1:1000; Abcam 113686;), mouse anti-yeast Aac2 (6H8; 1:1000; [73]) and Starbright 520/700-conjugated (BioRad) secondary antibodies.

### Statistical Analyses

Immunoblots and TLC plates were quantitated by Quantity One or ImageLab (BioRad Laboratories). Statistical comparisons (ns, *P >* 0.05; 1 symbol *P ≤* 0.05; 2 symbols *P ≤* 0.01; 3 symbols *P ≤* 0.001; 4 symbols *P ≤* 0.0001) were performed using Prism 11 (GraphPad). All graphs show the mean ± SD. The statistical tests executed and sample sizes are explicitly indicated in the associated figure legends.

### Miscellaneous

With the exception of the subcellular fractionation immunoblots, which are representative of two independent experiments performed on two separate days, all presented immunoblots and TLC images are representative of at least three independent experiments performed on three separate days.

## Acknowledgements

We would like to thank Drs. Mike Renne (Saarland University, Germany) for SD yeast media formulations, Carla Koehler (UCLA, USA) and Andy Reichert (Heinrich Heine University Düsseldorf, Germany) for antibodies, and Hiromi Sesaki (JHMI, USA) for reagents and equipment access. This manuscript is the result of funding in part by the National Institutes of Health (NIH; NIH grants R01GM151746, R01GM111548, R01GM111548-08S1, and R01GM111548-08S2 to SMC). It is subject to the NIH Public Access Policy. Through acceptance of this federal funding, NIH has been given a right to make this manuscript publicly available in PubMed Central upon the Official Date of Publication, as defined by NIH.

## Figure Legends

**Figure S1.** OM-Psd1 short-circuits the need for Ups2 or Mic60. Mitochondrial phospholipids were labeled overnight with ^14^C-acetate in indicated yeast grown in rich (A) dextrose or (G) lactate medium and resolved by TLC (different representative TLC images versus those shown in Figure 2 E and F are provided for each). Quantitation of mitochondrial PC:PE ratio (B, H), PI (C, I), PA (D, J), CL (E, K) amounts and PE:CL ratio (F, L) (mean ± SD for *n* = 6 biological replicates from 2 clones/genotype). Significant differences compared to the respective Ups2- and Mic60-proficient parent (^) or IM-Psd1 (+) were determined by one-way ANOVA with Tukey’s multiple comparisons.

