## Supplementary figures and images for "Rewiring Mitochondrial Phosphatidylethanolamine Metabolism Identifies New and Unaccounted Trafficking Steps"

### Supplemental Figure S1

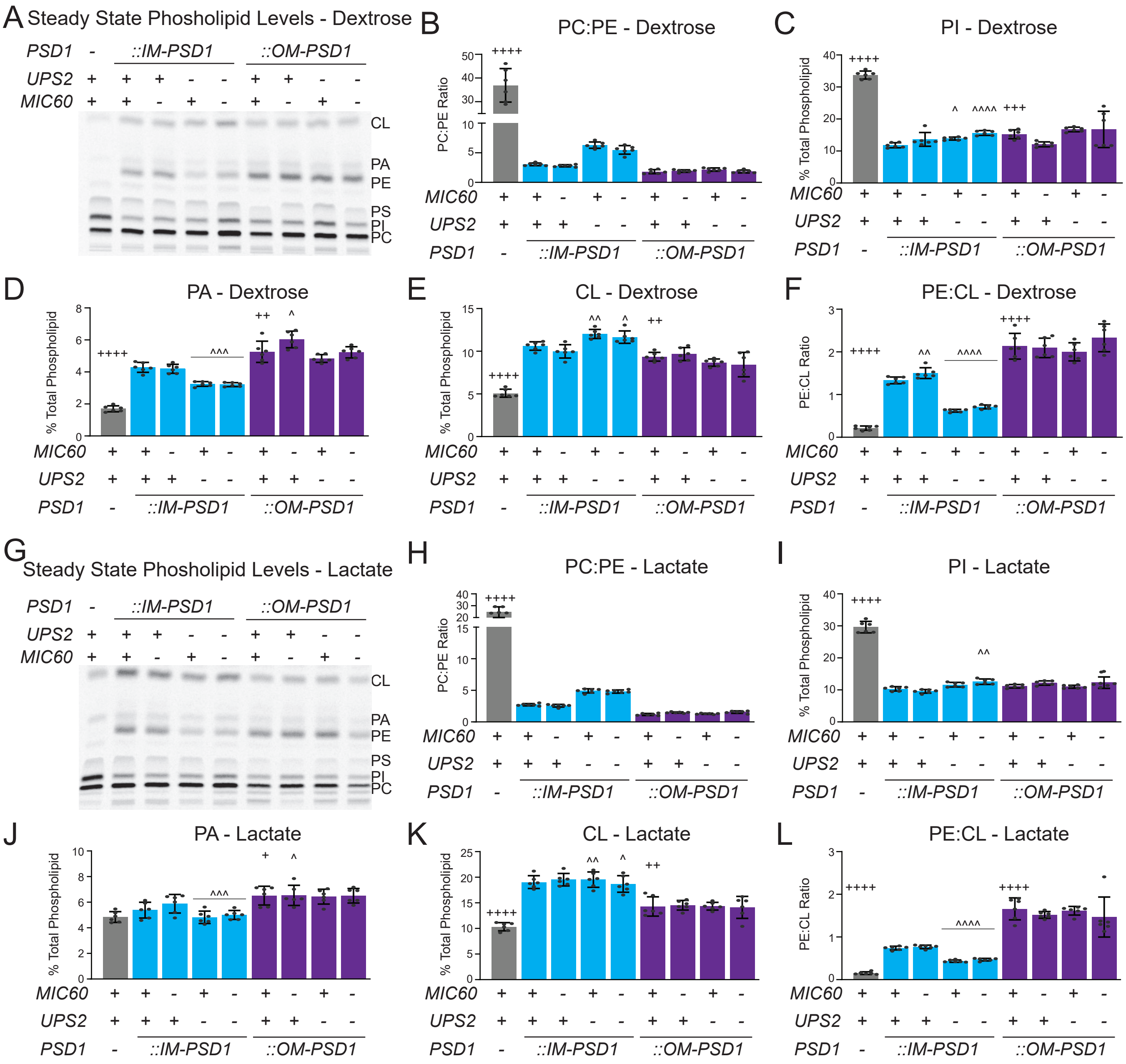


^ = vs IMPsd1; * = vs parent

Figure S1
